# *In vivo* CGG repeat RNA binding protein capture identifies RAN translation modifiers and suppressors of repeat toxicity

**DOI:** 10.1101/2021.01.08.425998

**Authors:** Indranil Malik, Yi-Ju Tseng, Shannon E. Wright, Kristina Zheng, Prithika Ramaiyer, Katelyn M. Green, Peter K. Todd

## Abstract

Fragile X-associated tremor/ataxia syndrome (FXTAS) is a neurodegenerative disorder caused by a transcribed CGG repeat expansion in the 5’ UTR of *FMR1*. Expanded CGG repeat RNAs both sequester RNA-binding proteins (RBPs) into nuclear foci and undergo repeat-associated non-AUG (RAN) translation into toxic homopolymeric peptides. RBPs that interact with CGG repeats may play a pivotal role in foci formation and/or RAN translation. Here we employed a CGG repeat RNA-tagging system to capture and identify CGG repeat binding RBPs *in vivo* under different cellular conditions. We found that several SR (serine/arginine-rich domain) proteins interact with CGG repeat RNAs basally and under cellular stress. These same proteins strongly modify toxicity in a Drosophila model of FXTAS, improving eye degeneration and survival. Furthermore, genetic or pharmacological targeting of the serine/arginine protein kinases (SRPKs) suppresses RAN translation in cellular reporters and toxicity in fly models of FXTAS and C9orf72 ALS/FTD. Finally, pharmacological targeting of SRPK1 supressed CGG repeat toxicity and enhanced survival in rodent neurons. Taken together, these findings demonstrate roles for CGG repeat RNA binding proteins in both RAN translation and repeat toxicity and suggest SRPK inhibition may serve as a possible therapeutic strategy in repeat expansion disorders.

## Introduction

Fragile X-associated Tremor/Ataxia Syndrome (FXTAS) is an age-related neurodegenerative disorder which impacts approximately 1/5000 people (Jacquemont *et al*, 2004; Tassone *et al*, 2012). It results from a transcribed CGG repeat expansion in the 5’UTR of *FMR1* (Hagerman *et al*, 2001). Patients develop imbalance, dementia, parkinsonism and tremors starting in their 50’s or 60’s (Jacquemont *et al*, 2003). Pathologically, the condition is associated with diffuse neuronal loss and brain atrophy as well as accumulation of ubiquitinated inclusions within neurons and glia throughout the brain (Greco *et al*, 2002, 2006; Ariza *et al*, 2016). FXTAS is an inexorably progressive and fatal condition without effective treatment. Thus identifying targetable factor (s) involved in CGG repeat expansion associated toxicity may help in develpoment of new therapeutics.

CGG repeats are thought to elicit toxicity through two non-exclusive mechanisms (Glineburg *et al*, 2018). Expanded repeat RNAs can elicit gain of function (GOF) toxicity by sequestering essential RNA-binding proteins (RBPs) and forming RNA-RNA and RNA-protein complex condensates (Jazurek *et al*, 2016; Jain & Vale, 2017; Glineburg *et al*, 2018). This pathologic process is best exemplified by the sequestration of muscleblind (MBNL) proteins by CUG repeat RNA in myotonic dystrophy type 1 (DM1) (Taneja *et al*, 1995; Miller *et al*, 2000). The repeat mRNA and MBNL protein avidly co-localize into RNA foci in both patient tissues and in model systems (Miller *et al*, 2000). Moreover, DM1 patients have splicing defects and clinical phenotypes that mimic those seen with genetic ablation of MBNL1 and upregulation of MBNL1 suppresses relevant disease phenotypes in DM1 disease models (Mankodi *et al*, 2000; Kanadia *et al*, 2003, 2006; Wang *et al*, 2019). At CGG repeat expansions that cause FXTAS, *in vitro* RNA pull down assays identified Pur alpha, hnRNP A2/B1 Sam68, and DROSHA/DGCR8 as potential repeat RNA targets (Jin *et al*, 2007; Sofola *et al*, 2007; Sellier *et al*, 2010, 2013). Overexpression of some of these factors in *Drosophila* can suppress CGG repeat elicited phenotypes (Jin *et al*, 2007; Sofola *et al*, 2007). However, direct manipulation of these factors has not yet recapitulated (in their absence) or suppressed (in their overexpression) disease relevant phenotypes in rodent or human neuronal model systems.

Repeat RNAs also support translational initiation in the absence of AUG start codons through repeat-associated non-AUG (RAN) translation (Zu *et al*, 2011). RAN translation is supported by many repeat expansions including CGG repeats associated with FXTAS and GGGGCC repeats associated with ALS/FTD (Ash *et al*, 2013; Mori *et al*, 2013; Todd *et al*, 2013; Bañez-Coronel *et al*, 2015; Zu *et al*, 2017; Soragni *et al*, 2018; Ishiguro *et al*, 2017). RAN translation occurs across multiple reading frames of the same repeat to produce toxic repetitive polypeptides that accumulate in patient neurons and tissues (Zu *et al*, 2013; Krans *et al*, 2016; Gendron *et al*, 2013; Zu *et al*, 2017; Soragni *et al*, 2018). Expression of dipeptide repeat products resulting from C9 ALS/FTD GGGGCC RAN translation are sufficient to elicit toxicity in model systems even in the absence of repetitive RNA, suggesting that they play an active role in disease pathogenesis (Mizielinska *et al*, 2014; Wen *et al*, 2014; Jovičić *et al*, 2015; May *et al*, 2014; Lee *et al*, 2016; Zhang *et al*, 2018, 2019).

In FXTAS, RAN translation in the GGC reading frame generates a polyglycine containing peptide termed FMRpolyG, which accumulates into ubiquitinated inclusions in both model systems and patient tissues (Todd *et al*, 2013). Induced expression of repeats that support FMRpolyG synthesis elicit toxicity in heterologous cells, rodent neurons, flies and transgenic mice (Todd *et al*, 2013; Sellier *et al*, 2017; Buijsen *et al*, 2014). Moreover, sequence manipulations that suppress RAN translation of FMRpolyG largely preclude CGG repeat associated toxicity in overexpression systems (Todd *et al*, 2013). RAN translation is selectively activated by cellular stress response pathways that typically preclude translational initiation, suggesting that specific translational factors or alternative mechanisms may underlie RAN translation and its contributions to repeat associated toxicity (Green *et al*, 2017; Cheng *et al*, 2018; Sonobe *et al*, 2018; Westergard *et al*, 2019). Indeed, unbiased and targeted genetic approaches have identified potential factors that preferentially modulate RAN translation including ribosomal protein RPS25 and RNA helicase DDX3X (Linsalata *et al*, 2019; Yamada *et al*, 2019; Cheng *et al*, 2019).

One potential confounder from studies of repeat RNA binding proteins in FXTAS and other repeat expansion disorders to date is their reliance on *in vitro* repeat RNA capture methodologies (Jazurek *et al*, 2016). As most RNAs come to interact with specific RBPs during transcription, export and/or translation as part of their normal life cycle in the cell, we reasoned that critical factors involved in both RAN translation and RNA foci formation/RBP sequestration might be missed with *in vitro* assays, which do not capture these interactions with great fidelity. It is also possible that interactions of specific factors with CGG repeat RNA may only occur in particular cellular states, such as after activation of cellular stress pathways. Dynamic RNA-protein and RNA-RNA interactions change under cellular stress (Van Treeck & Parker, 2018; Matheny *et al*, 2020) and repeat RNAs such as ALS/FTD-associated C9ORF72 GGGGCC repeats can partition into stress granules and interact with specific proteins (Fay *et al*, 2017). Thus, capturing context-specific repeat RNA-protein interactions *in vivo* might reveal novel modulators of repeat RNA biology and pathogenesis.

To define the roles played by CGG repeat RNA binding proteins in both RAN translation and FXTAS pathogenesis *in vivo*, we developed a repeat RNA-tagging system, which allows for the unbiased identification of repeat RNA-binding proteins inside cells (Harlen & Churchman, 2017). We fused a pathogenic CGG repeat expansion containing reporter with PP7 viral stem loops, which bind to viral coat protein with high affinity (Chao *et al*, 2008). By co-expressing the coat binding protein, PCP, we were able to isolate CGG repeat RNAs and associated RBPs. This modular system can be utilized in the context of cellular perturbations such as stress induction or drug treatment. Using this technique, we found that multiple SR proteins interact with CGG repeat RNAs under normal and stress conditions. Genetic targeting of serine/arginine proteins suppresses rough eye phenotypes and extends survival in a *Drosophila* model of FXTAS. Moreover, genetic or chemical targeting of serine/arginine protein kinases (SRPK1) that regulate SRSF1 function selectively suppress RAN translation and toxicity in fly and rodent neuronal models of FXTAS. Taken together, these data present a novel approach to identify repeat RNA-binding proteins *in vivo* and establish SR protein kinases as a possible target to modulate RAN translation and repeat expansion-associated toxicity.

## Results

### Development of a CGG repeat RNA-tagging system

Cellular RNA-binding proteins (RBPs) play critical roles in RNA gain-of-function toxicity in repeat expansion disorders. To identify RBPs that interact with expanded CGG repeat RNAs that may modulate RAN translation in a FXTAS disease model, we designed an RNA-tagging system that allowed isolation and identification of CGG repeat RNA-binding proteins inside cells (Harlen & Churchman, 2017). To accomplish this, we modified previously characterized CGG RAN translation-specific nanoluciferase (nLuc) reporters by inserting PP7 viral stem-loops after the stop codon (**Fig 1A**) (Kearse *et al*, 2016). This reporter system allows translation of the CGG RAN reporter, while keeping the PP7 stem-loop structures unperturbed to interact with the PP7 coat protein (PCP). This construct was co-transfected with PP7 coat-binding protein containing a nuclear localization signal and a 3xFLAG epitope tag (PCP-NLS-3xFLAG), which facilitated immunoprecipitation (IP) using anti-FLAG antibody (**Figure 1A**). Consistent with previous findings, CGG RAN translation efficiency was significantly less than AUG-driven canonical translation, but it was enhanced with activation of integrated stress response by thapsigargin (TG) treatment (**Figure 1B-C**) (Green *et al*, 2017).

**FIGURE 1.**
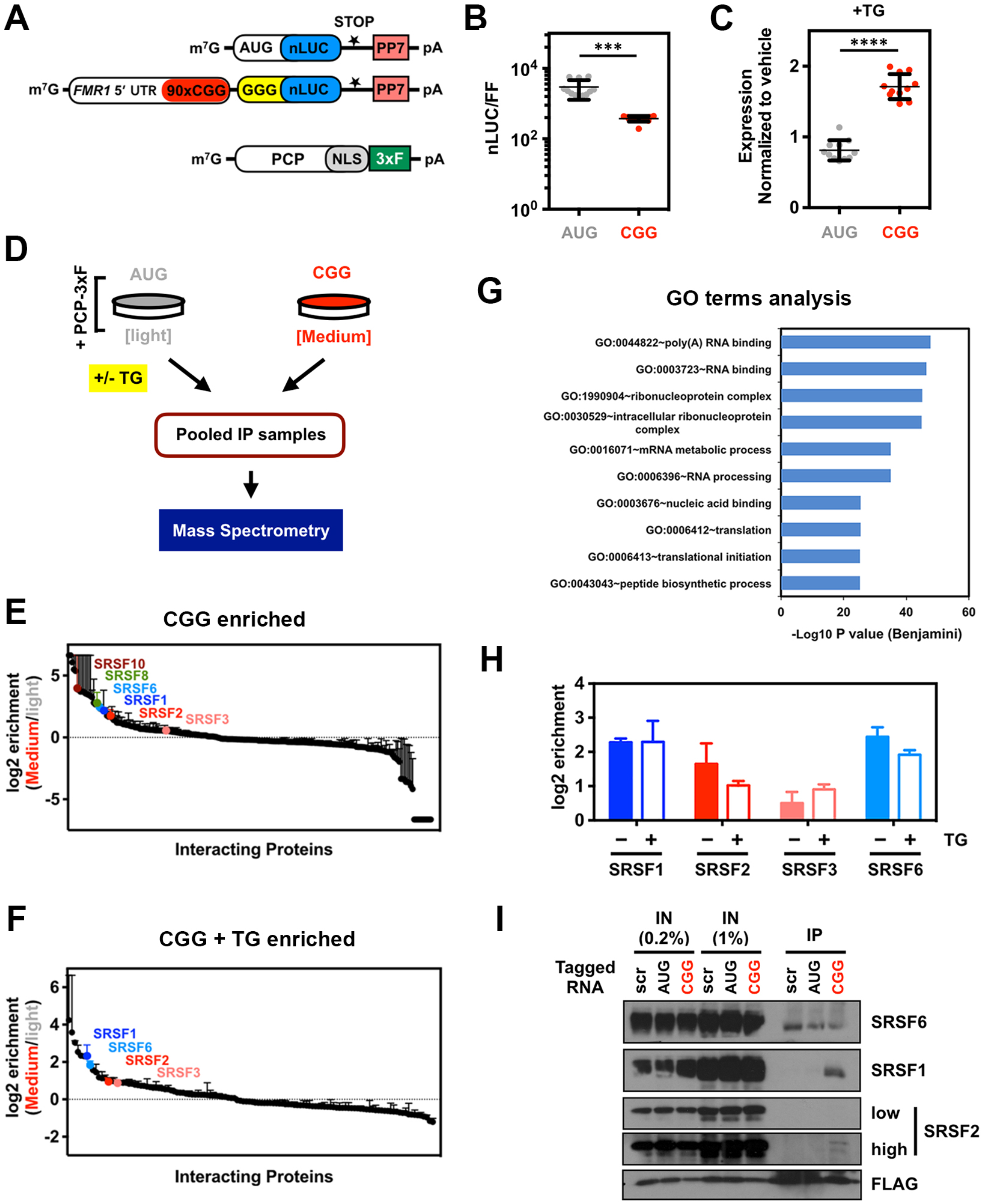
Repeat RNA-tagging system enables identification of CGG repeat specific RNA-binding proteins within cells. (A) Schematic of PP7-tagged RNA reporters and PCP-NLS-FLAG constructs used in this study. (B) Relative expression from PP7-tagged CGG-nLuc reporters compared to AUG-driven reporters in HEK293T cells (n=9). (C) Expression of AUG-control and CGG-nLuc RAN translation reporters in HEK293T cells treated with the ER stress agent thapsigargin (TG, 2 μM) (n=9) normalized to vehicle (DMSO). (D) Schematic of immunoprecipitation and mass spectrometry experiments aimed at identifying CGG repeat RNA interacting proteins (see Methods for details). (E-F) Log_2_ fold-change of the CGG-interacting protein enrichment compared AUG-reporter (n=2 independent experiments; error bars represent range between repeats) under normal (E) and after integrated stress response activation by TG (F). (G) GO term analysis of manually curated differentially enriched CGG-interacting proteins. (H) Log_2_ fold-change of SRSF proteins: 1, 2, 3 and 6 in CGG-reporter enrichment compared to AUG-reporter basally and after TG treatment (n=2 independent experiment; error bars represent range between two repeats). (I) Co-immunoprecipitation of indicated SRSF proteins with scrambled (scr), AUG and CGG-tagged reporter constructs. SRSF1 and 2 specifically immunoprecipitated with CGG-tagged reporter. For graphs in (B-C) error bar represents +/− SD. Statistical analysis was performed using: Two-tailed Student’s t test with Welch’s correction, ***p < 0.001; ****p < 0.0001.

To quantitatively identify ribonucleoprotein complexes formed by the CGG-nLuc reporter, we used SILAC (stable isotope labeling by amino acids in cell culture) with HEK293T cells grown in light and medium amino acids before IP (**Figure 1D**) (Ong *et al*, 2002). We used an AUG-nLuc–PP7 as control (Figure 1A) and a GGGGCC-nLuc-PP7 for comaparative analysis (see Methods and for details). For this study we only focus on the CGG-interactome, as the GGGGCC-interactome will be reported in a separate manuscript (Extended data Table). AUG-nLUC-PP7 reporter was grown in light and CGG- nLUC-PP7 reporter was grown in medium amino acids containing HEK293T cells and both of them were co-transfected with the 3xFLAG tagged PCP. In parallel, to determine RBPs that may interact with these reporters after ISR activation, cells were treated with 2uM TG 5 hours before the IP. The SILAC analysis provided more than 300 protein interactors with quantification of their binding preferences for AUG and CGG reporters (**Figure 1E**, **Supplementary Figure 1 and Extended data Table**). Fewer interactors were identified after ISR activation with TG treatment (**Figure 1F** and **Supplementary Figure 1**). A reduction in RNA-protein interaction during integrated stress response might be a result of perturbation in normal RNA metabolism during cellular stress (Bond, 2006). Gene ontology (GO) enrichment analysis of manually curated CGG-enriched protein interactors using the Database for Annotation, Visualization and Integrated Discovery (DAVID) yielded top functional categories related to poly(A) RNA-binding, RNA-binding, ribonucleoprotein complexes, mRNA metabolic processes, RNA processing and translation (**Figure 1G**), indicating that this RNA-tagging IP successfully captured RBPs that may differentially interact with CGG repeat RNA during its synthesis, transport and translation (Dennis *et al*, 2003).

### SR proteins selectively interact with CGG repeat RNAs

Both AUG initiated and CGG repeat constructs formed similar ribonucleoprotein complexes and exhibited significant overlap in their interactomes (**Supplementary 1A-B**). However, several RBPs including multiple serine/arginine-rich domain (SR) proteins preferentially interacted with CGG repeat reporters compared to the AUG reporter (**Figure 1E-F; Supplementary Figure 1C-D**). Top RBPs that preferentially interact with CGG repeat reporter in basal condition (no stress) include heterogeneous nuclear ribonucleoproteins hnRNPH, hnRNPC; poly(a) binding protein PABPN1, PABPC1 and PABPC4; ribosomal proteins RPL18A, RPLP0 and RPLP2 (**Supplementary Figure 1C**). Stress specific interactors include ribosomal protein L40 (UBA56), hnRNPQ, PABPC1, LARP1 and YBX1 (**Supplementary Figure 1D)**. Interestingly, multiple SR proteins interact with the CGG reporter both basaly and in presence of stress. In particular, SRSF 1, 2, 3 and 6 showed enrichment both basally and in response to cellular stress (**Figure 1H**). SR proteins are a large family of RBPs consisting 12 structurally related proteins containing characteristic Arg/Ser-rich (RS) domains that influence mRNA splicing, export, stability and translation (Zhou & Fu, 2013). To validate these interactions, we immunoprecipitated CGG RNA-tagged reporter RNA and immunoblotted for SRSF proteins. SRSF1 interacted very strongly with CGG RNA-tagged reporter compared to a scrambled RNA-tag or AUG-reporter (**Figure 1I**). SRSF2 also interacted preferentially with the CGG RNA-tagged reporter, while SRSF6 interacted non-specifically with all mRNAs (**Figure 1I**).

### SR proteins modulate CGG repeat RNA toxicity

To test if any of the CGG repeat RNA interactors can modulate CGG RNA toxicity, we conducted a candidate-based screen using a *Drosophila melanogaster* model of FXTAS (**Figure 2A**) (Todd *et al*, 2013; Linsalata *et al*, 2019). This fly model carries an upstream activation sequence (UAS)-driven 5’ UTR of human *FMR1* with 90 CGG repeats fused to EGFP in the +1 (FMRpolyG) reading frame. Expression of this reporter in the fly eye via a GMR-GAL4 driver results in production of ubiquitin-positive aggregates of the RAN translated FMRpolyG, leading to a rough-eye phenotype (Todd *et al*, 2013). We have previously used this fly model to screen for modifiers of *FMR1* CGG RAN translation and identified several translation-associated factors that modulate CGG repeat toxicity (Linsalata *et al*, 2019). For the modifier screen in this study, we selected a few candidates from the list of top CGG-interacting proteins (**Supplementary Figure 1C-D**) to cross with GMR-GAL4 control and 90 CGG repeats expressing FXTAS model flies. Candidate modifiers with intrinsic toxicity were excluded from further analysis (**Supplementary Figure 2A**). We found that knocking down *Drosophila* homologs of several SR proteins (SRSF1, 2/8, 4/6 and 7/3) as well as translation initiation factor eIF3G, ribosomal protein RPLP0 and RNA helicase DHX30 significantly reduced CGG repeat RNA toxicity without eliciting significant toxicity in isolation (**Figure 2B-C**; **Supplementary Figure 2B**). Factors that significantly enhance CGG repeat toxicity in fly eye include fly homologs of heterogeneous nuclear ribonucleoproteins hnRNP H/F and hnRNPQ; poly(a) binding protein (PABPN1), nuclear cap-binding protein (NCBP1) and DExD-Box Helicase 39B (**Figure 2B**).

**FIGURE 2.**
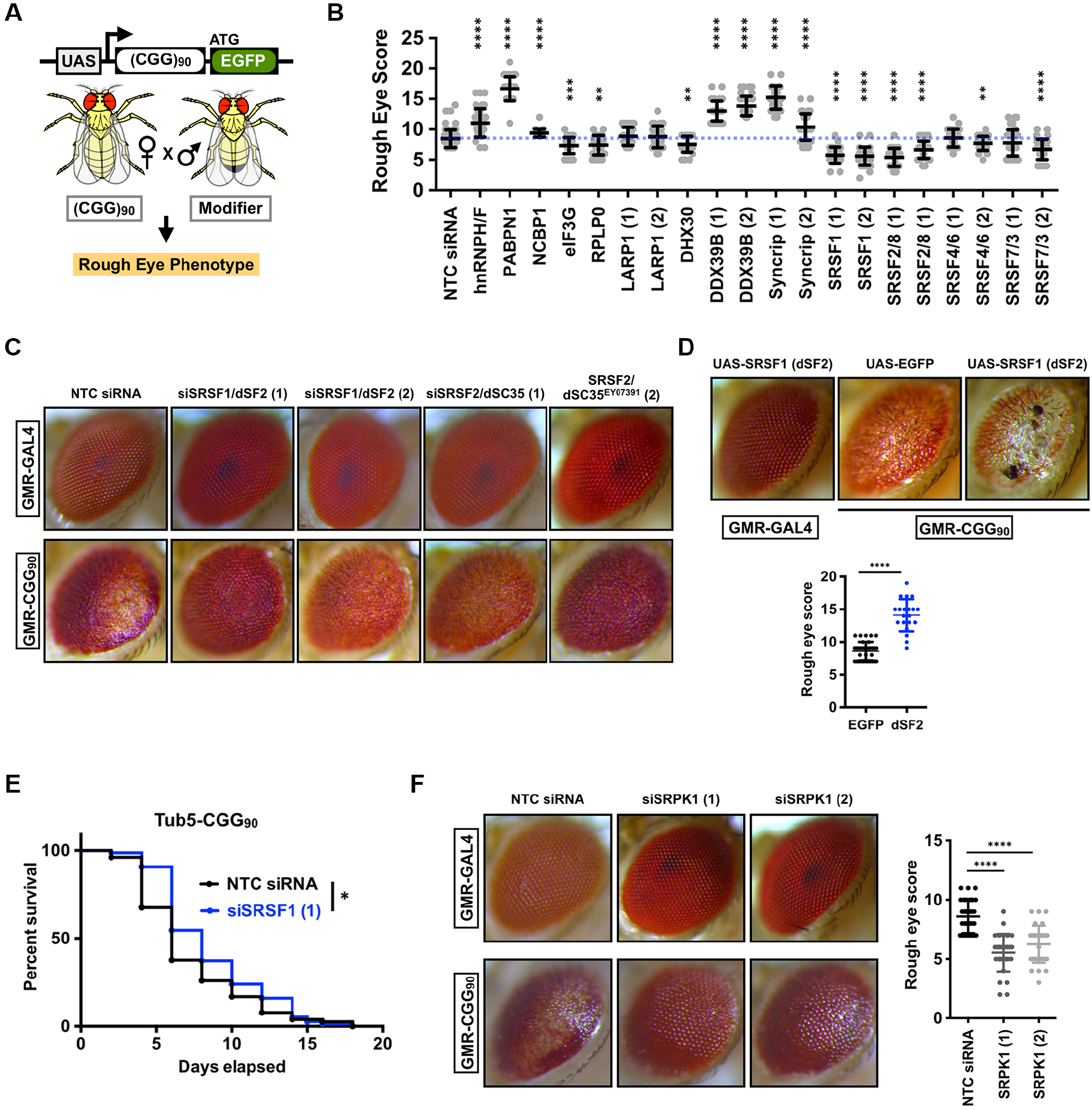
SRSF proteins act as modifiers of CGG repeat RNA associated toxicity in *Drosophila*. (A) Schematic of (CGG)90-EGFP construct and experimental outline for rough eye phenotype. (B) Quantitation of GMR-GAL4 driven uas-(CGG)90-EGFP eye phenotype with candidate modifiers (t-test with Welch corrections for comparisons with the control; n ≥ 30 flies/genotype). **p < 0.01; ***p < 0.001; ****p < 0.0001. [NTC = non targeting control] (C) Representative photographs of fly eyes expressing either GMR-GAL4 driver alone or with uas- (CGG)90-EGFP construct, with fly SRSF1 (dSF2) and SRSF2 (dSC35) knockdown or disruptions. (D) Representative photographs of fly eyes as in (C) and quantitation as in (B) with SRSF1/dSF2 overexpression (dSF2 OE). (E) Survival assay on flies expressing (CGG)90-EGFP under Tub5-GS driver with drug initiation starting 1 day post eclosion and continuing through experiment (Log-rank Mantel–Cox test; n = 78-80/genotype) with SRSF1 knockdown *p < 0.05. (F) Representative photographs of fly eyes as in (C) and quantitation as in (B) with siRNA mediated knockdown of SRPK1 (dSRPK1). Details of fly genotypes described in supplementary Table 2.

To investigate whether overexpression of SR proteins might influence CGG repeat toxicity, we developed a transgenic *Drosophila* model carrying UAS-driven dSF2 (*Drosophila* homolog of human SRSF1) and expressed this reporter in the fly eye using a GMR-GAL4 driver. Expression of dSF2 alone did not lead to any eye abnormalities. However, co-expression of dSF2 significantly enhanced CGG repeat elicited eye degeneration compared to an eGFP control (**Figure 2D**). Ubiquitous expression of 90 CGG repeats after eclosion in adult flies using the Tub5-Geneswitch system significantly shortens lifespan (Todd *et al*, 2010; Linsalata *et al*, 2019). Expression of siRNAs against SF2/SRSF1 under an inducible Tub5-Geneswitch driver significantly increased the lifespan of flies expressing 90 CGG repeats (**Figure 2E**). Similarly, SF2/SRSF1 increased lifespan when CGG repeat was expressed selectively within neurons in adult flies under an inducible Geneswitch ElaV driver (**Supplementary 3A**). However, siRNAs against SRSF2/SRSF8 failed to enhance survival of flies expressing 90 CGG repeats (**Supplementary FIgure 3B**). These results suggest that SF2/SRSF1 in particular plays a role in adult onset CGG repeat associated neurodegeneration in *Drosophila.*

Serine-arginine protein kinases (SRPKs1-3) selectively phosphorylate SR proteins and modulate their subcellular localization and play active roles in spliceosome assembly (Zhou & Fu, 2013). In addition, many non-splicing functions for SPRKs have been reported, including roles in tau phosphorylation and Alzheimer’s disease (AD) pathogenesis (Hong *et al*, 2012). SRPKs exhibit highly tissue-specific expression profiles, suggesting that these kinases may have specialized functions (Nakagawa *et al*, 2005; Wang *et al*, 1998). Since multiple SR proteins modulate CGG repeat RNA toxicity in flies, we tested the effects of altering expression of SRPK1 on CGG repeat RNA toxicity in our FXTAS fly model. Interestingly, we found that selective knockdown of the *Drosophila* homolog of SRPK1 (dSRPK1) reduced CGG repeat RNA toxicity and significantly improved the CGG repeat rough eye phenotype (**Figure 2F**). However, expression of siRNAs against dSRPK1 ubiquitously under an inducible Tub5 Geneswitch driver did not enhance lifespan of flies expressing 90 CGG repeats (**Supplementary Figure 3C**), perhaps because of some basal toxicity.

### SR proteins modify GGGGCC repeat RNA toxicity in *Drosophila*

Recent studies have shown similarities in mechanisms underlying RAN translation initiation at CGG and GGGGCC repeats (Green *et al*, 2017; Glineburg *et al*, 2018; Kearse *et al*, 2016). Furthermore, ‘GC-rich’ sequences in both these repeats may also exhibit similar GOF toxicity through ‘RNA-gelation’ and sequestration of related RBPs (Jain & Vale, 2017; Glineburg *et al*, 2018). Indeed, a recent study has shown that SRSF1 is required for nuclear export of GGGGCC repeats and knockdown of SRSF1 modulates GGGGCC repeat toxicity in a fly model (Hautbergue *et al*, 2017). Thus, we asked if SR proteins that interact with CGG repeats and rescued CGG repeat toxicity in our fly screen may modulate GGGGCC repeat toxicity in a *Drosophila* model. To this end, we used a transgenic Drosophila model expressing UAS-driven 28 GGGGCC repeats that exhibits a rough-eye phenotype when expressed in fly eyes (He *et al*, 2020). We found that *Drosophila* homologs of several SR proteins, including SF2/SRSF1, SRSF2/8 and 4/6, significantly reduced rough-eye phenotypes in the GGGGCC repeat expressing fly (**Figure 3A-B**). Rough-eye phenotypes of this fly are enhanced several fold at higher temperatures (29°C), presumably due to enhanced transgene expression (**Figure 3C**). Enhanced phenotypes include presence of severe necrosis and shrinkage of the total eye. Expression of siRNAs against SF2/SRSF1 was able to significantly reduce necrosis and prohibit shrinkage of GGGGCC repeat expressing fly eyes (**Figure 3D-E**). Next, we tested if SR proteins modulate GGGGCC repeat elicited shortening of lifespan (He *et al*, 2020). As with CGG repeats, ubiquitous expression of siRNA against SRSF1 significantly enhanced survival of GGGGCC repeat expressing fly (**Figure 3F**). Moreover, expression of siRNA against SRPK1 (dSRPK1) reduced GGGGCC repeat toxicity and significantly improved the associated rough eye phenotype (**Figure 3G-H**). Additionally, unlike CGG repeat expressing fly, ubiquitous expression of siRNAs against dSRPK1 under an inducible Tub5-Geneswitch driver significantly enhanced lifespan of flies expressing GGGGCC repeats (**Supplementary Figure 3E**).

**FIGURE 3.**
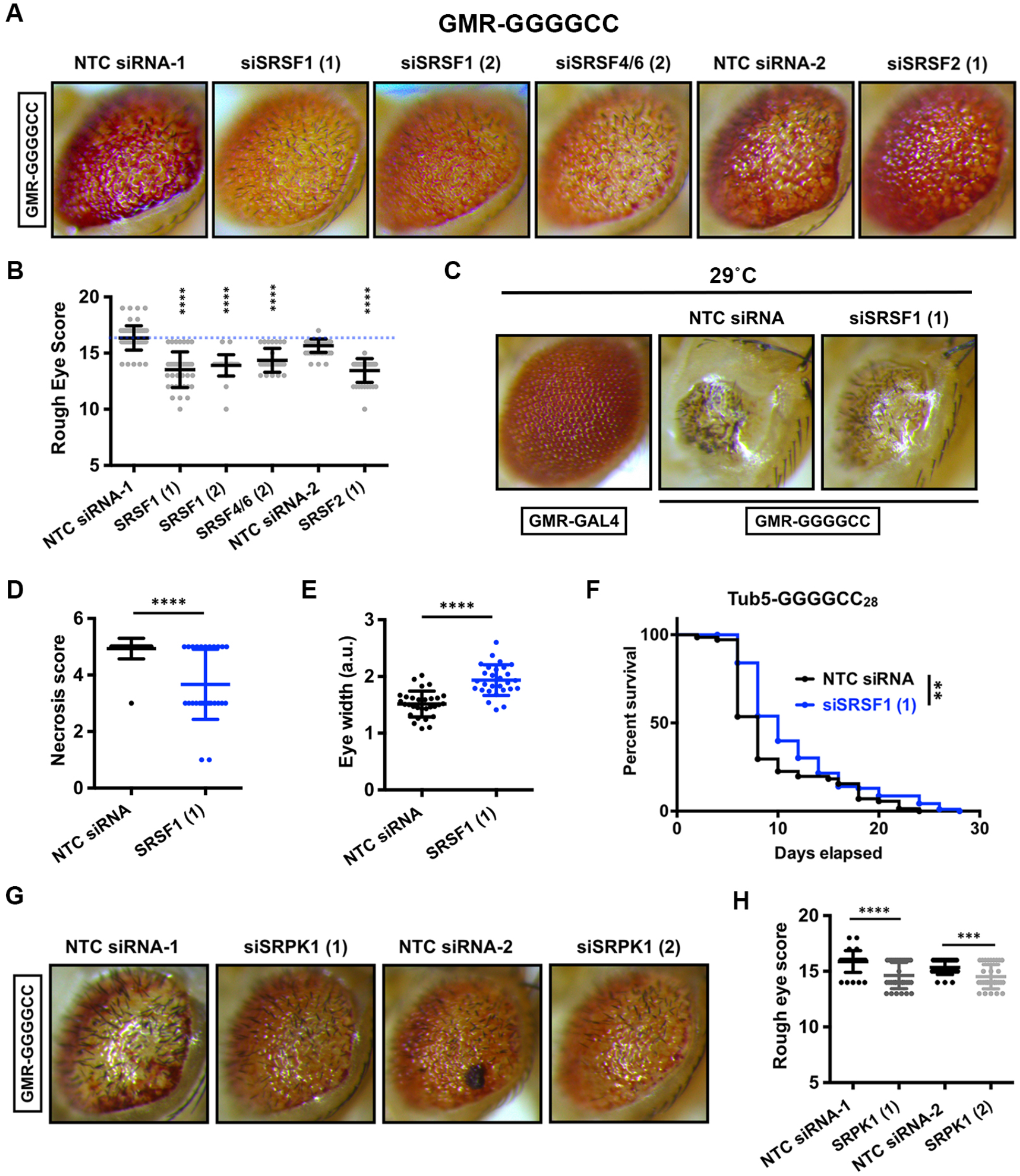
SRSF proteins modify GGGGCC repeat RNA-associated toxicity in a Drosophila model. (A) Representative photographs of fly eyes expressing GMR-GAL4 driven (GGGGCC)28-EGFP with indicated uas-siRNAs to fly SRSF proteins at 25°C. (B) Quantitation of GMR-GAL4 driven (GGGGCC)28-EGFP eye phenotype with SRSF modifiers (t-test with Welch corrections for comparisons with the control; n ≥ 30 flies/ genotype). ****p < 0.0001 (C) Representative photographs of fly eyes expressing either GMR-GAL4 driven (GGGGCC)28-EGFP at 29°C with siRNA against SRSF1. Quantification of eye necrosis (D) and eye diameter (E) at 29°C in GMR-GAL4 driven (GGGGCC)28-EGFP flies. (F) Survival assays of (GGGGCC)28-EGFP expressing fly under Tub5-GS driver (Log-rank Mantel–Cox test; n = 71-90/genotype) with control or SRSF1 siRNA. **p < 0.01 (G) Representative photographs of fly eyes expressing GMR-GAL4 driven (GGGGCC)28-EGFP with siRNA mediated knockdown of SRPK1 or disruption by insertion. (H) Quantitation of rough eye phenotypes.. t-test with Welch corrections for comparisons with the control; n ≥ 30 flies/ genotype). ***p < 0.001; ****p < 0.0001

### Loss of SRSF1 alters CGG RNA distribution and enhances CGG RAN translation

SR proteins including SRSF1 play critical roles in mRNA export and influence the stability and translation of target transcripts. Consistent with this, a previous study has found that SRSF1 is required for NXF1-dependent nuclear export of GGGGCC repeat RNA and knockdown of SRSF1 significantly reduces RAN translated products from a GGGGCC reporter (Hautbergue *et al*, 2017). Thus, we next asked whether SR proteins that interact with CGG repeats might also play a similar role in facilitating CGG RAN translation. We used our previously characterized nLuc-based CGG RAN translation reporter consisting of a 3xFLAG-tagged nanoluciferase (nLuc-3XF) downstream of the 5’ UTR of human *FMR1* (**Figure 4A**) (Kearse *et al*, 2016). These reporters can be used for detection of RAN products by either luminescence based assays or Western blotting. As opposed to the prior study, we observed that knockdown of SRSF1 led to a modest enhancement in CGG RAN translation as detected by both luciferase assay and Western blotting (**Figure 4B-C**). We confirmed this observation with an independent siRNA targeting SRSF1 (**Supplementary Figure 4B-C**). Next, we tested if other SRSFs that successfully rescued fly phenotype can modulate RAN translation. While the knockdown of SRSF2 significantly inhibited CGG RAN translation, it also had a significant effect on AUG-driven control translation. Indicating these observed effects may not be RAN-translation specific (**Supplementary Figure 4A**). Knockdown of another SR protein, SRSF8, inhibited AUG-driven translation, but has no significant effect on CGG RAN translation. Thus, the impacts of genetic modulation of SR proteins on RAN translation were inconsistent and not directly correlative with their impacts on toxicity in *Drosophila.*

**FIGURE 4.**
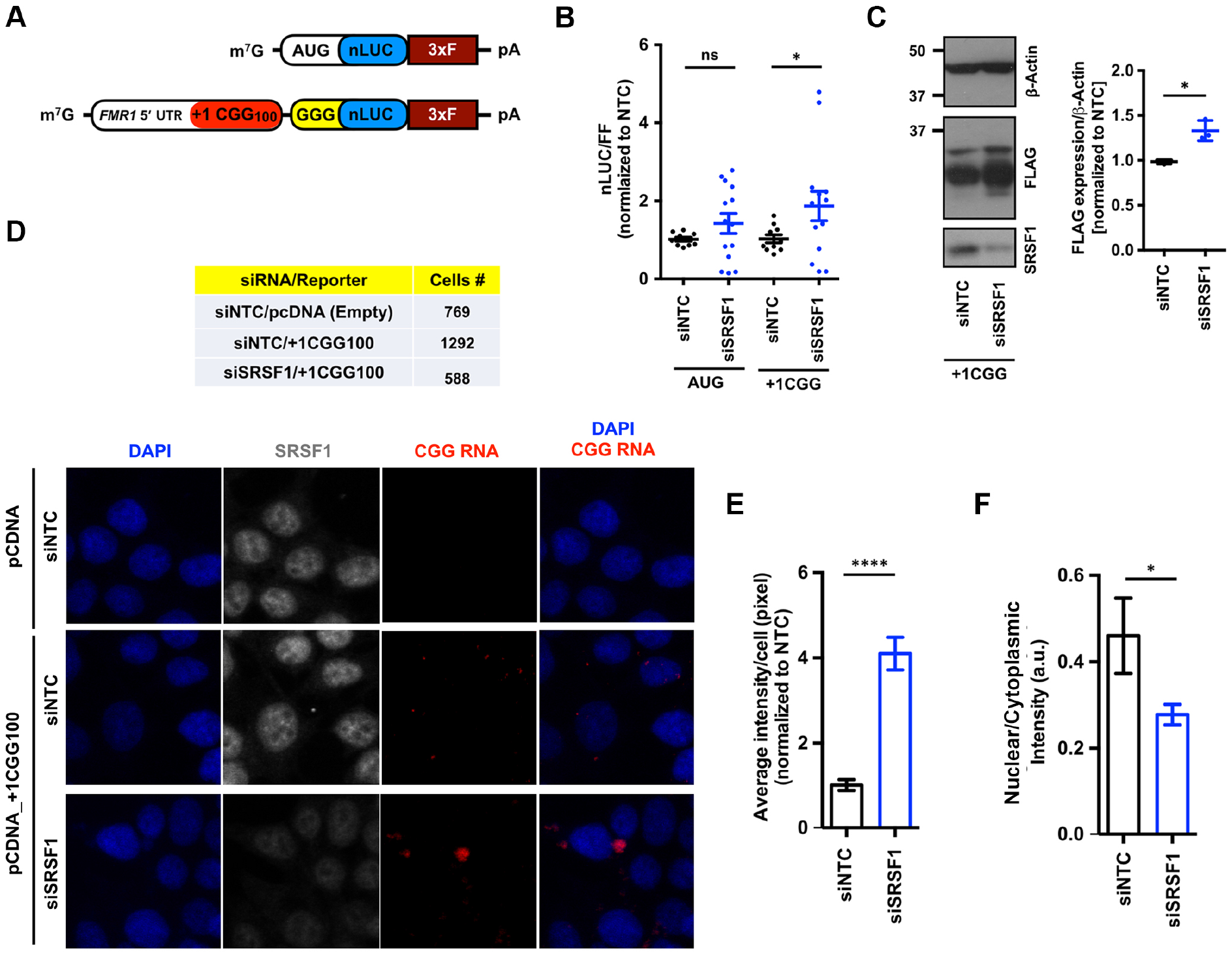
SRSF1 Knockdown altered CGG RNA localization and enhanced CGG RAN translation. (A) Schematic of +1CGG RAN translation nLuc-3xFLAG and AUG-driven control nLuc-3xFLAG reporters. (B) Relative expression of AUG-nLuc and CGG-nLuc reporters in HEK293T cells (n=10-14) following knockdown of SRSF1. Comparisons between siEGFP and siSRSF1 treated cells. (C) Anti-FLAG Western blot of FMRpolyG nLUC-3xFLAG with and without SRSF1 knockdown in HEK293T cells (n=3). Error bars represent mean +/−SEM. t-test with Welch corrections. *p < 0.05 (D) Representative images of subcellular distribution of CGG-nLuc-3xFLAG reporter RNAs using HCR with and without SRSF1 knockdown in HEK293T cells. n= # of cells used for quantification for each condition as mentioned in the table. Quantification of CGG-nLuc-3xFLAG reporter RNA intensity (E) and nuclear/cytoplasmic distribution (F). Error bars represent mean +/−SEM. t-test with Bonferroni and Welch’s correction *p < 0.05, ****p < 0.0001.

The RNA binding protein MBNL1 binds to and regulates the subcellular distribution of CCUG repeat RNAs, which in turn influences RAN translation of these repeats in reporter systems (Zu *et al*, 2017). Given that SRSF1 knockdown enhanced CGG RAN translation, we evaluated whether SRSF1 knockdown might alter the subcellular distribution of CGG repeat reporter RNAs. To test this, we used Hybridization Chain Reaction (HCR) to detect the CGG RNA along with immunocytochemistry (ICC) to detect SRSF1 in HEK293T cells (Rodriguez *et al*, 2020; Glineburg *et al*, 2021). In HEK293T cells expressing the CGG-nLuc RAN reporter, we observed both nuclear and cytoplasmic RNA foci (**Figure 4D**). Knockdown of SRSF1 led to an significant increase in CGG repeat RNA foci size and intensity, perhaps due to increased stability of the reporter RNA (**Figure 4D-E**). Moreover, reduction of SRSF1 led to enhanced cytoplasmic distribution of CGG reporter RNA (**Figure 4D and F**). Together these results suggest that nuclear SRSF1 proteins interact with expressed CGG repeats and retain them in the nucleus. This reduces their cytoplasmic transit and accumulation and thus suppresses their translation. Following SRSF1 knockdown, the repeat RNAs fail to form nuclear foci and move more readily to the cytoplasm where they undergo RAN translation. These data suggest that loss of SRSF1 suppresses CGG repeat RNA toxicity primarily through a suppression of nuclear CGG repeat RNA elicited toxicity, which can be quite robust in isolation (Sellier *et al*, 2010).

### SRSF Protein Kinase 1 (SRPK1) inhibitors modulate CGG RAN translation

Since genetic ablation of *Drosophila* homolog of SRPK1 strongly modified rough eye phenotypes in both CGG and GGGGCC expressing flies, but knockdown of select SR proteins had differential effects on RAN translation, we next asked what impact SRPK1 inhibition might have on RAN translation. In order to test the effects of SRPK1 inhibition on RAN translation, we used two known chemical inhibitors of SRPK1 (**Figure 5A**). First, we tested the effects of SRPIN340, an ATP-competitive SRPK inhibitor, on RAN translation by pre-treating HEK293T cells with SRPIN340 followed by transfecting CGG RAN translation reporters (Fukuhara *et al*, 2006). SRPIN340 treatment at 50 uM led to a significant and selective decrease in +1CGG RAN translation as detected by western blot (**Figure 5B**). Similarly, SRPIN340 treatment inhibited CGG RAN translation as measured by nanoLuciferase assay, but this had no impact on AUG-nLuc translation (**Figure 5C**). This effect was not isolated to FMRpolyG synthesis, as SRPK1 inhibition by SRPIN340 also suppressed GGGGCC RAN translation (GA70) and +2CGG RAN translation (FMRpolyA) using our previously published reporters (**Figure 5D**). As pharmacological agents may have off-target effects, we also evaluated the impact of a second SRPK inhibitor, SPHINX31 (Gammons *et al*, 2013; Batson *et al*, 2017). Similar to SRPIN340, SPHINX31 significantly and selectively inhibited +1CGG, GGGGCC and +2CGG RAN translation (**Figure 5E-F**). These results indicate that pharmacological inhibition of SRPK1 has a general inhibitory effect on RAN translation. RAN protein levels increase under various stress conditions, including ER stress (Green *et al*, 2017). As SPRK inhibitors selectively suppressed RAN translation, we wondered if SRPK inhibition might also impede stress-induced enhancement of RAN translation. Pre-treating cells with 50 uM SRPIN340 led to a complete blockade of thapsigargin-induced enhancement of +1CGG RAN ranslation as detected by western blotting and luciferase assays (**Figure 6A-B**). SRPIN340 also suppressed stress-induced GGGGCC (GA70) and +2CGG RAN translation (**Figure 6B-D**). Together these results suggested that pharmacological inhibition of SRPK1 can inhibit both basal level and stress-induced increase in RAN translation across at least two different repeats and at least two reading frames of the CGG repeat.

**FIGURE 5.**
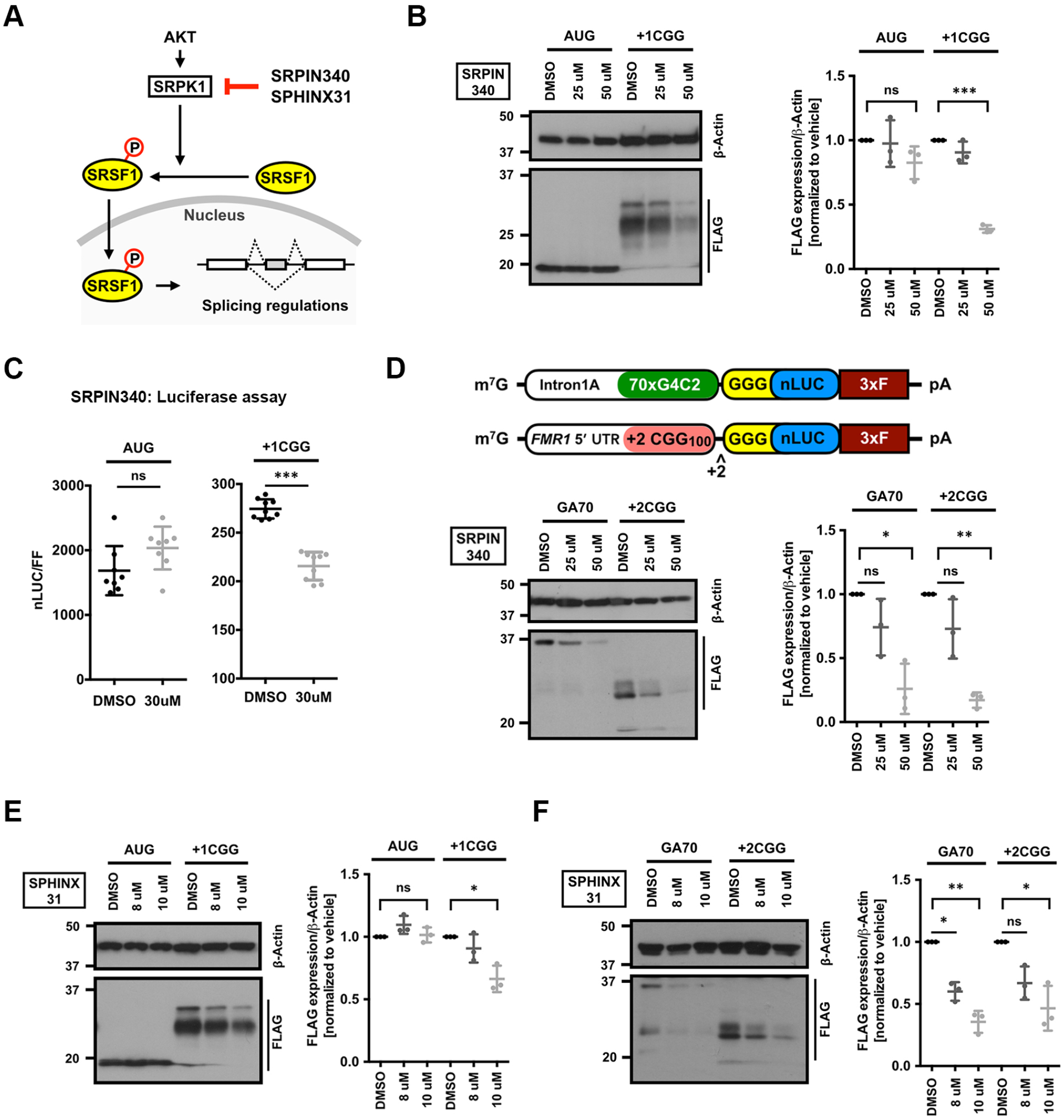
SRPK1 inhibitors selectively suppress RAN translation. (A) Schematic of AKT/SRPK1 signaling pathway that regulate SRSF1 localization with example of known pharmacological compounds that inhibit SRPK1. (B) Anti-FLAG Western blot of DMSO and SRPIN340 pre-treated HEK293T cells expressing AUG-nLUC-3xFLAG control or CGG-nLuc-3xFLAG RAN translation reporters. β-Actin is used as a loading control. To prevent signal saturation, AUG-nLuc lysate was diluted 1:3 in sample buffer prior to loading (n=3). (C) Relative expression of AUG-nLuc and CGG-nLuc reporters in HEK293T cells (n=8-9) following treatment with DMSO and SRPIN340. (D) Anti-FLAG Western blot of DMSO and SRPIN340 pre-treated HEK293T cells expressing GGGGCC-nLuc-3xFLAG (GA70) and +2CGG-nLuc-3xFLAG (FMRpolyA) RAN translation reporters (n=3). Schematics of the GA70 (GGGGCCx70) and +2CGG reporters presented on top. (E) Western blot of DMSO and SPHINX31 pre-treated HEK293T cells expressing AUG-nLUC-3xFLAG control or CGG-nLuc-3xFLAG RAN translation reporters. (n=3). (F) Anti-FLAG Western blot of DMSO and SPHINX31 pre-treated HEK293T cells expressing GGGGCC-nLuc-3xFLAG (GA70) and +2CGG-nLuc-3xFLAG (FMRpolyA) RAN translation reporters (n=3). Error bars represent mean +/− SD. *p < 0.05; **p < 0.01 and ***p < 0.001. To prevent over-exposure, the AUG-nLuc lysate was diluted 1:3 in the sample buffer.

**FIGURE 6.**
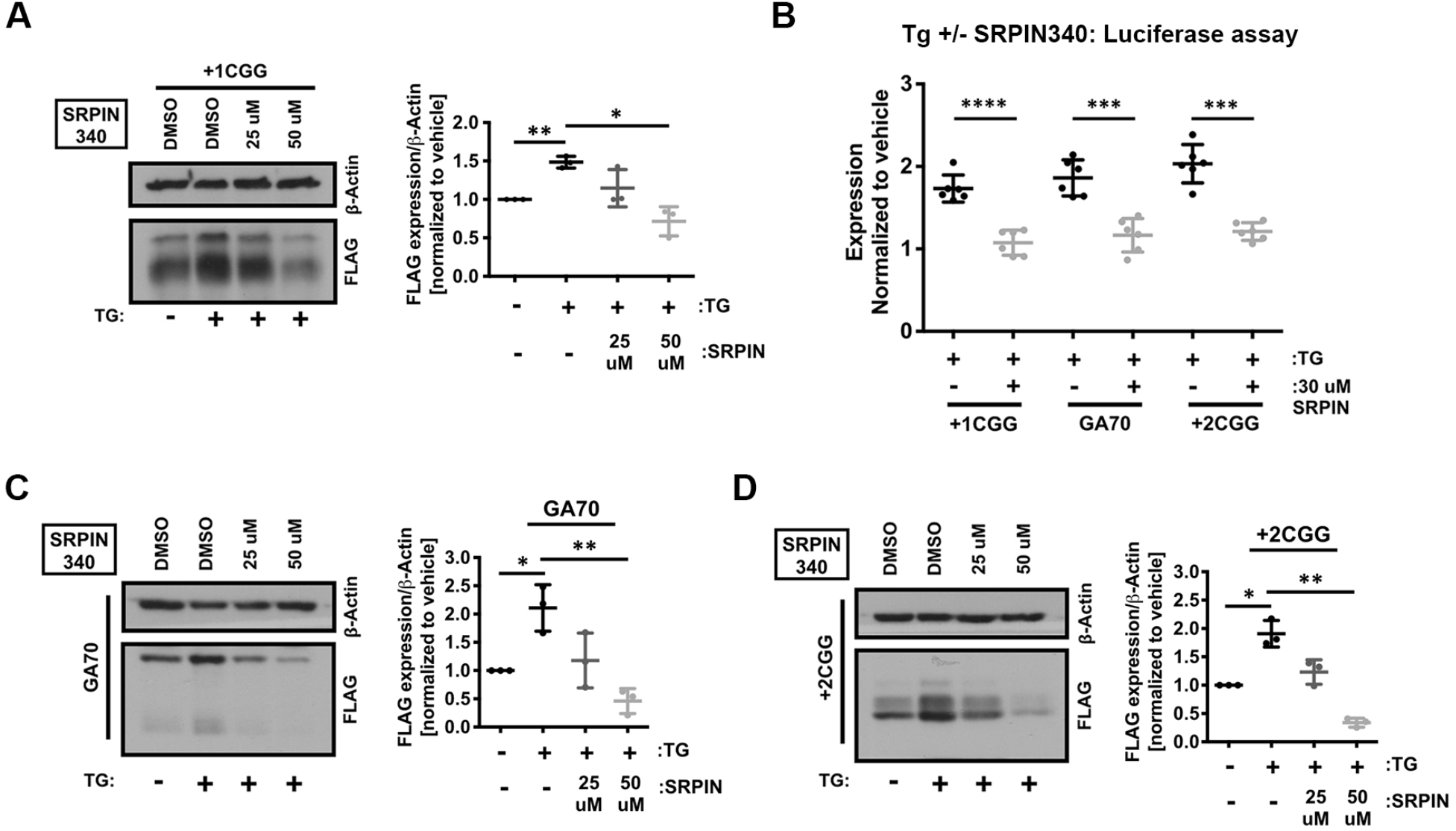
SRPK1 inhibition prevents stress-induced enhancement of RAN translation. (A) Expression of +1CGG-nLuc-3xFLAG RAN translation reporters in HEK293T cells treated with 2 μM TG (for stress induction) analyzed by Western blotting (n=3). To evaluate effects of SRPK1 inhibition cells were pre-treated with DMSO or SRPIN340 before reporter transfection. (B) Relative expression of +1CGG-nLuc reporters in HEK293T cells (n=6) following stress induction with 2 μM TG treatment. Values normalized to vehicle (DMSO) treatment. As in (A), cells were pre-treated with DMSO or SRPIN340 before reporter transfection. Expression of GGGGCC-nLuc-3xFLAG (C) and +2CGG-nLuc-3xFLAG (D) RAN translation reporters in HEK293T cells treated with 2 μM TG (for stress induction) analyzed by Western blotting (n=3). To evaluate effects of SRPK1 inhibition cells were pre-treated with DMSO or SRPIN340 before reporter transfection. Error bars represent mean +/− SD. *p < 0.05; **p < 0.01 and ***p < 0.001

### SRPK1 inhibition enhances survival in (CGG)100 expressing neurons

As pharmacological inhibition of SRPK1 suppressed RAN translation from both +1 FMRpoly(G) and +2 FMRpoly(A) translation frames in mammalian cells and knockdown of SRPK1 in Drosophila also suppressed CGG repeat toxicity, we asked whether SRPK1 inhibitors might mitigate CGG repeat toxicity in mammalian neurons. To this end, we expressed a +1(CGG)_100_-EGFP reporter encoding for +1 FMRpoly(G) in rat primary neurons along with an mApple reporter plasmid that allowed for selective tracking of transfected cells using an automated fluorescence microscopy assay system (Linsalata *et al*, 2019; Barmada *et al*, 2015). We used an AUG-driven EGFP reporter as a control for transfection and exogenous protein expression-associated toxicity. Consistent with previous results(Linsalata *et al*, 2019), we observed that +1(CGG)_100_-EGFP expression markedly reduced neuronal survival compared to EGFP expression, as tracked by automated longitudinal fluorescence microscopy over 10 days (**Figure 7A-B, Supplementary Figure 5A-D**). Next, we treated neurons with SRPIN340 at a range of concentrations from 10 to 50 uM before +1(CGG)_100_-EGFP reporter transfection (**Supplementary Figure 5A-B**). Neuronal survival rate was significantly improved with SRPIN340 treatment at concentrations of 30, 40 and 50 uM compared to DMSO treatment. However, SRPIN340 appeared to have some neurotoxicity itself on EGFP transfected neurons at higher concentrations. SRPIN340 treatment at 40 uM showed the most favorable and selective effects on neuronal survival **(Figure 7A, Supplementary Figure 5A-B**). To confirm that this suppression of neurotoxicity by SRPIN340 treatment is not a drug-specific effect, we also tested the ability of SPHINX31 to suppress +1(CGG)_100_-EGFP induced toxicity. Similar to SRPIN340, SPHINX31 significantly improved +1(CGG)_100_-EGFP expressing neuronal survival at all tested concentrations (**Supplementary Figure 5C-D**). SPHINX31 elicited maximum suppression of neurotoxicity at concentrations of 8 and 10 uM, consistent with our observation of inhibition of RAN translation in a similar range of concentrations. At 8 uM concentration, SPHINX31 showed a promising effect on neuronal survival against +1(CGG)_100_-EGFP reporter-induced toxicity over any intrinsic drug associated toxicity (**Figure 7B**). Together, these findings suggest that pharmacological inhibition of SRPK1 can suppress neurotoxicity of expanded CGG repeats through inhibiting RAN translation.

**FIGURE 7.**
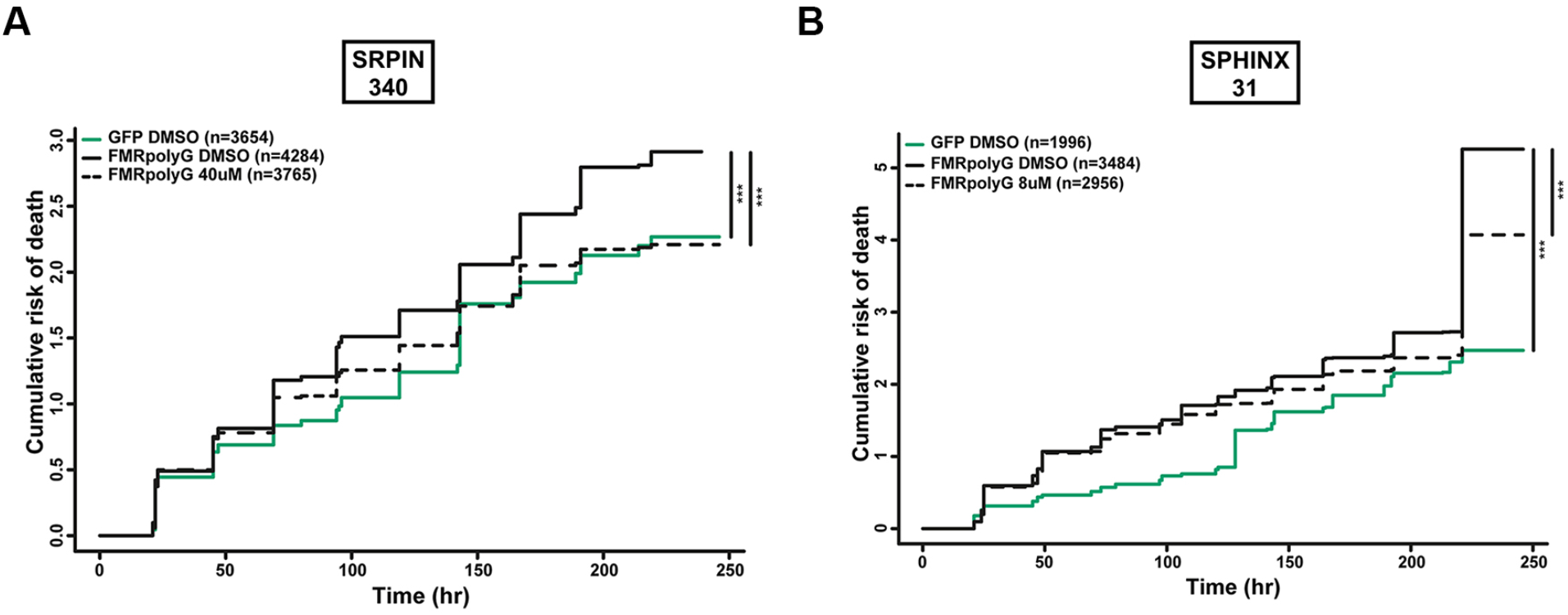
SRPK1 inhibition mitigates (CGG)100 RNA toxicity in primary rat cortical neurons. Pharmacological targeting of SPRK1 with (A) 40 uM SRPIN340 and (B) 8 uM SPHINX31 reduced the cumulative risk of death in +1(CGG)100-EGFP expressing neurons (Cox proportional hazard analysis). (n= # of neurons quantified for each conditions as mentioned within the graphs); ***p < 0.001

## Discussion

Nucleotide repeat expansions as RNA form *in vivo* structures that complex with specific RNA binding proteins within different cellular compartments (Krzyzosiak *et al*, 2012; Jazurek *et al*, 2016; Ciesiolka *et al*, 2017). These interactions influence their toxicity and have the potential to alter their stability, distribution and translation. To date, most studies of repeat RNA-protein interactions have relied on *in vitro* capture methods that do not take the cellular context of the mRNA into consideration (Jazurek *et al*, 2016). As such, they have the potential to miss important interacting proteins that might be of lower overall abundance or require interaction within the context of either specific cellular compartments or *in vivo* RNA structures. Here we utilized an alternative method for identifying RNA-RBP interactors that allows for capture of complexes that form within cells under both basal conditions and in response to cellular stress. By applying this tool to CGG repeats that cause FXTAS, we identified a series of novel interactors, including factors with differential interaction profiles after stress induction (**Figure 1E-F**). Evaluation of these interactors identified SRSF proteins as playing direct roles in CGG repeat RNA localization, RAN translation, and toxicity in model systems. Moreover, genetic and pharmacological targeting of the major SRSF kinase (SRPK1) significantly impaired RAN translation at multiple GC rich repeat sequences and suppressed toxicity in *Drosophila* and CGG repeat expressing rodent neurons (**Figure 5–7**). Taken together, these studies highlight the value of *in vivo* tools for identification of RNA-RBP interactions and suggest that SRPK may serve as a therapeutic target worthy of further evaluation in repeat expansion disorders.

Our screening identified multiple SR proteins that interact with CGG repeat RNA and modify CGG repeat toxicity (**Figure 1E-F and 2B**). Notably, SRSF1, 2, 3 and 6 interact with CGG repeats normally and under stress conditions (**Figure 1H**), and loss of SR protein orthologues in *Drosophila* suppressed CGG repeat toxicity (**Figure 2**). Though SR proteins play a major role in splicing, they are also implicated in mRNA export, regulation of RNA stability and translation (Jeong, 2017). The selective enrichment of SR proteins with CGG repeat-containing reporters that do not undergo splicing indicates that these factors may influence CGG repeat stability, distribution and translation. Consistent with this, we observed CGG repeat specific but somewhat discordant effects of knockdown of different SR proteins on both CGG RAN translation and its distribution within cells (**Figure 4**). SRSF1, 2 and 9 have been previously shown to interact with other repeat RNAs, with functional implications for GGGGCC repeat RNA associated with C9ALS (Sato *et al*, 2009; Donnelly *et al*, 2013; Lee *et al*, 2013; Cooper-Knock *et al*, 2014). While sense GGGGCC RNA has been shown to interact with both SRSF1 and SRSF2, antisense C4G2 RNA has been shown to colocalize with SRSF2 (Cooper-Knock *et al*, 2015). Interestingly, analysis of C9ALS patient cerebellum samples have shown extensive alternative splicing (AS) defect in transcripts targeted by hnRNPH1 and SRSF1, indicating that SRSF1-sequestration by GGGGCC RNA may account for loss-of-function toxicity (Prudencio *et al*, 2015; Conlon *et al*, 2016).

Although genetic ablation of SRSF1 mitigated rough eye phenotypes and extended survival in *Drosophila* model of FXTAS, knockdown of SRSF1 led to a modest increase in transiently transfected +1CGG reporter RAN translation (**Figure 2 and Figure 4B-C**). This increase in RAN translation correlates with enhanced cytoplasmic distribution of the reporter CGG RNAs (**Figure 4D-F**). This observation is consistent with a recent study finding that nuclear retention of CCUG repeat transcripts by MBNL1 inversely correlates with RAN translation (Zu *et al*, 2017). Knockdown of MBNL1 enhanced CCUG transcript export and enhanced RAN translation in reporter systems. However, this result is different than that observed for GGGGCC repeats, where a recent study found that SRSF1 depletion inhibits nuclear export of GGGGCC repeat RNA and reduced generation of RAN products, which allows for an alleviation of neurotoxicity in *Drosophila* (Hautbergue *et al*, 2017). We observed similar effects of SRSF1 depletion on rough eye phenotype and survival of a GGGGCC repeat expressing fly (**Figure 3A-F**). The exact cause of this discrepancy, however, is not clear. At least a portion of the explanation may result from the constructs used, where the CGG repeat here is embedded within a 5’UTR of an mRNA that may undergo uORF like non-AUG translation (Kearse *et al*, 2016; Rodriguez *et al*, 2020). As such, its default state may be cytoplasmic transport. In contrast the GGGGCC repeat constructs used in past studies may preferentially reside in the nucleus, such that loss of SRSF1 favors their retention. This notion is supported by the observation that knockdown of SRSF1 increases the overall intensity of CGG RNA signal as measured by HCR and the formation of large cytoplasmic RNA granules in SRSF1 depleted cells (**Figure 4E**).

An alternative explanation is that reduction of SRSF1 in *Drosophila* may alter other pathways, which indirectly lead to phenotypic correction. For example, the major alternative splicing target of SRSF1 is MKNK2, encoding for the Mnk2 enzyme. Mnk2 is one of the two serine/threonine kinases (Mnk1 and Mnk2) involved in both Ras-MAPK pathway and eukaryotic translation initiation factor 4E (eIF4E) phosphorylation (Karni *et al*, 2007; Anczuków *et al*, 2012). It is known that SRSF1 overexpression modulates alternative splicing of *MKNK2*, to reduce the longer isoform Mnk2a and increase the shorter isoform Mnk2b (Maimon *et al*, 2014). While SRSF1 knockdown has an opposite effect on alternative splicing of *MKNK2*. The Mnk2b isoform lacks a C-terminal MAPK-binding domain and cannot activate the p38-MAPK required for stress-induced cell death (Maimon *et al*, 2014). Thus, it is possible that knockdown of SRSF1 enhances the Mnk2a isoform leading to the activation of p38-MAPK and overall increase in survival.

Besides SRSFs, our dual screens identified a number of other intriguing RNA-binding proteins that may play a role in CGG repeat associated toxicity. Some of these proteins mitigate rough eye phenotypes in flies, such as DHX30 and eIF3G, while others such as hnRNPQ enhance rough eye phenotypes (**Figure 2B**). DHX30 is an ATP-dependent DEAD/H RNA helicase that has been implicated in translation regulation apoptosis associated transcripts (Rizzotto *et al*, 2020). A previous *Drosophila* screen identified the fly homolog of DHX30 as a modest modifier of rough eye phenotype in this FXTAS fly model, which we also observed here (Linsalata *et al*, 2019). Given the roles of other DEAD box helicases in RAN translation, including DDX3, the selective identification of this factors as a CGG repeat RNA interactor suggests the need for further study of how it might alter repeat RNA behavior (Linsalata *et al*, 2019). Similarly the mammalian eIF3 complex protein eIF3F has already been implicated in RAN translation at CAG and GGGGCC repeats (Ayhan *et al*, 2018). While eIF3G is one of the core subunits, eIF3F a non-core regulatory subunit of the eIF3 complex (Hinnebusch, 2006). Identification of this complex in association with the repeat RNA suggests that it may play a role in its translation, especially given prior studies suggesting eIF3 may be critical in non-canonical initiation events. Lastly, hnRNPQ is implicated as a key factor in controlling the translation of FMRP from FMR1 transcripts through an IRES mediated mechanism (Choi *et al*, 2019). Given recent studies linking CGG repeats, RAN translation and FMRP synthesis, this factor will require further evaluation in RAN translation assays at this and other repeat structures.

Protein kinases are attractive targets for drug development. Notably, SPRK1 inhibitors are actively pursued as potential anti-cancer drugs (van Roosmalen *et al*, 2015; Chandra *et al*, 2020). SRPK1 phosphorylates multiple serine residues in the RS1 domain of SRSF1 to regulate its nuclear localization (Zhou & Fu, 2013). Altered levels of SRSF1 have been reported in many cancers, where phosphorylation of SRSF1 plays a decisive role in alternative splicing of disease associated transcripts (Anczuków *et al*, 2015; Sheng *et al*, 2018). Furthermore, in some cancers misregulation of SRPK1 has been linked with cell proliferation, migration, and angiogenesis (van Roosmalen *et al*, 2015). In light of these findings our discovery of SRPIN340 and SPHINX31 as potential inhibitors of RAN translation is potentially worthy of further pursuit. However, there may be some intrinsic toxicity associated with these compounds, given that we observed modest toxicity in neurons expressing GFP alone, especially for SRPIN340 (**Supplementary Figure 5**). Interstingly, SPHINX31, which has been shown to inhibit SRPK1 more potently than SRPIN340, appeared relatively less toxic indicating it to be a more potenital candidate for future studies (**Supplementary Figure 5**) (Gammons *et al*, 2013). Finally, since knowckdown of SRSF1 didn’t have an inhibitory effect on +1CGG RAN translain, it is possible SRPK1 inhibitors are not acting directly through SRSF1. There are other possiblities such as indrect effects on SRSF1-associated signalling pathways as decribed earlier. It is also possible that at high concentrations SRPK1 inhibitors effect both SRPK1 and SRPK2, thus affecting multiple SRSF proteins simultaneous leading to an cumulative effect of RAN translation. Futrher studies are needed to decipher the mechanims by which SRPK1 inhibitors inhibit RAN translation.

In sum,we have developed an in vivo method for identifying Repeat RNA binding proteins and applied it to CGG repeats to reveal both novel interactors and phenotypic modifiers associated with these repeat expansions. Future comparative studies with other repeat elements and in other systems should allow for a better understanding of Repeat RNA-protein complex formation and interactions in vivo and guide us in our understanding of the native functions of repeat elements, and how repeats as RNA cause disease and potentially what drug targets are likley to serve as effective therapeutics in these currently untreatable disorders.

## Methods

### 1. Antibodies

For western blots following antibodies were used: FLAG-M2 at 1:1000 dilution (mouse, Sigma F1804), 1:2500 β- Actin (mouse, Sigma A1978), 1:1000 SRSF1 (Rabbit, Proteintech 12929-2-AP), 1:1000 SRSF2 (Rabbit, Proteintech 20371-1-AP), 1:1000 SRSF6 (Rabbit, Bethyl Laboratories A303-669A-T) in 5% non-fat dry milk. HRP-conjugated goat-anti-mouse (115-035-146) or goat-anti-rabbit (111-035-144) secondary antibodies (Jackson ImmunoResearch Laboratories) were used at a 1:10,000 dilution in 5% non-fat dry milk. For HCR: - Primary Antibodies for ICC - SRSF1 1:100, rabbit from Proteintech, 12929-2-AP; Secondary antibodies for ICC - 1:500, Alexa Fluor 488 goat anti-rabbit IgG from Thermo Fisher Scientific, A11008.

### 2. Plasmids

Nano Luciferase (nLuc) reporters cloned into pcDNA3.1(+) vector encoding AUG-nLuc-3xFLAG, +1CGG100-nLuc-3xFLAG (FMRpolyG), +2CGG100-nLuc-3xFLAG (FMRpolyA), GGGGCC70-nLuc-3xFLAG (GA70) sequences used in this paper are published earlier (Green *et al*, 2017; Linsalata *et al*, 2019; Kearse *et al*, 2016). 2x PP7 stem loop sequence as described earlier (Harlen & Churchman, 2017; Coulon *et al*, 2014) was synthesized from GeneWitz. In order to make the RNA-tagging construct pcDNA3.1(+) AUG-nLuc-3xFLAG and pcDNA3.1(+) +1CGG100-nLuc-3xFLAG vectors were modified in two steps. First, nLuc reporter sequence was PCR modified to introduce a stop codon after nLuc sequence and remove the 3xFLAG sequence. This PCR product was cloned back in to the original vectors to make pcDNA3.1(+) AUG-nLuc and pcDNA3.1(+) +1CGG100-nLuc vectors. Finally, 2x PP7 stem loop were cloned into these vectors using ApaI restriction site to finally make - pcDNA3.1(+) AUG-nLuc-PP7 and pcDNA3.1(+) +1CGG100-nLuc-PP7 constructs. PCP-NLS sequence was PCR amplified from a PCP containing plasmid to introduce SV40 NLS sequence right after PCP. The PCP sequence-containg plasmid was generously gifted by Brittany Flores (Flores *et al*, 2019) and originally reported here (Yan *et al*, 2016). Then a 3xFLAG sequence was amplified from the AUG-nLuc-3xFLAG plasmid and PCR sewed with the PCP-NLS fragment to clone in pcDNA3.1(+) vector using KpnI and ApaI restriction sites to finally make the pcDNA3.1(+) PCP-NLS-3xFLAG construct.

See Supplementary **Table 1** for reporter sequences.

### 3. Cell culture, drug treatments and reporter assays

HEK293T cells were purchased from American Type Culture Collection (ATCC) and cultured in DMEM media supplemented with 10% FBS. They were confirmed to be mycoplasma free. Luminescence assays were performed as described earlier with slight modifications (Green *et al*, 2017; Linsalata *et al*, 2019). Briefly, HEK293T cells were seeded in 96-well plates at a concentration of 2 × 104 cells/well and transfected ~24 h later at ~ 70% confluency with 25 ng nLuc reporter plasmid and 25 ng pGL4.13 Firefly luciferase reporter plasmid using a ratio of 2:1 jetPRIME (Polyplus) to DNA following manufacturer’s recommendation. ~24 h post transfection cells were lysed with 70 uL Glo Lysis buffer (Promega) by incubating for 5 min on a shaker at room temperature. Then, 25 uL of lysate was mixed with NanoGlo substrate diluted 1:50 in NanoGlo buffer (Promega) and 25 uL of lysate was mixed with ONE-Glo luciferase assay buffer (Promega) in opaque 96-well plates. Reaction was allowed to continue for 5 min on a shaker in the dark. Finally, luminescence was measured on a GloMax 96 Microplate Luminometer.

For SRPK1 inhibitors, HEK293T cells were plated as described above and pre-treated with SRPIN340 (Sigma 5042930001) and SPHINX31 (BioVision, B2516-5) at desired concentrations for 8 h and 6 h before the transfection, respectively. Transfections and luminescence assays were performed as described above. For luminescence assays following ISR activation, HEK293T cells were seeded and transfected as described before for 19 h, followed by 5 h of treatment with 2 uM Thapsigargin (Thermo Fisher Scientific). All drugs were dissolved in DMSO and stored as recommended by the manufacturer.

For western blots, HEK293T cells were seeded in 24-well plates at a concentration of 1.5×105 cells/mL and transfected 24 h later at ~70% confluency with 250 ng nLuc reporters using a ratio of 2:1 jetPRIME as described earlier. 24 h post transfection cells were lysed in 300 uL of RIPA buffer containing protease inhibitor cocktail (cOmplete™ Mini, Sigma) for 30 min at 4°C with occasional vortexing. Lysates were cleared by centrifugation at 14000 rpm for 10 min, mixed with 6x SDS sample buffer and boiled at 90°C for 10 min before running on SDS-PAGE. If required lysates were stored at −80 °C for future experiments. For western blotting after drug treatment or stress induction, HEK293T cells were seeded in 24-well plates at 1.5×105 cells/mL and treatments were performed as described earlier. For each experimental condition at least three biological samples were run on 10% SDS-PAGE along with a standard curve for quantification of protein expression. Band intensities were measured using ImageJ and plotted using GraphPad Prism.

### 4. Tagged RNA capture and mass spectrometry

For tagged-RNA immunoprecipitaion (IP), HEK293T cells were grown in SILAC medium for 5–6 passages before transfection. DMEM deficient in L-arginine and L-lysine is supplemented with 10% dialyzed FBS and isotopes of lysine (final concentration 146 mg/liter) and arginine (final concentration 84 mg/L) for triplex SILAC labeling. For light SILAC DMEM, L-lysine (Lys-0) and L-arginine (Arg-0); for medium SILAC DMEM, L-lysine-4,4,5,5-d4 (Lys-4) and L-arginine [13C6] HCl (Arg-6); while for heavy medium L-lysine [13C6, 15N2]HCl (Lys-8) and L-arginine [13C6, 15N4]HCl (Arg-10) were added to the medium and filter-sterilized with a 0.2-μm filter. For transfection, cells were seeded at 2×106 cells/10 cm dish and transfected at ~70% confluency with 5.25 ug of PP7-tagged RNA bait plasmids [pcDNA3.1(+) AUG-nLuc-PP7, pcDNA3.1(+) +1CGG100-nLuc-PP7 and pcDNA3.1(+) (GGGGCC)70-nLuc-PP7] along with 0.75 ug of PCP-NLS-3xF plasmid, using jetPRIME reagents. Light SILAC was used for AUG-nLuc-PP7, medium SILAC +1CGG100-nLuc-PP7 and heavy SILAC was used for 1(+) (GGGGCC)70-nLuc-PP7 reporter. For ISR activation, cells were seeded similarly and transfected for 19 h, followed by 5 h of treatment with 2 uM Thapsigargin. Three 10 cm dishes were used per condition. 24 h after transfection cells were isolated by trypsinization and washing with 1xPBS. Cell pellets were immediately flash frozen using liquid nitrogen and proceeded to IP. Cell pellets from three 10 cm plates were pooled together and lysed in 1 mL of NP40 buffer (supplemented with cOmplete™ Mini protease inhibitor, 1mM PMSF, NEB Murine RNase inhibitor and RNaseIN) by incubating at 4 °C for 30 min with occasional pipetting to mix. Lysates were cleared by centrifugation at 20000xg for 10 min and the supernatant was transferred into a new tube. Protein concentration for each sample was measured by BCA assay (23227, Thermo Fisher Scientific). For IP, 3 mg of total protein was used for each lysate. Lysates were first incubated with 40 uL of pre-washed protein G beads for 30 min at 4°C to block any non-specific interaction. Then incubated with pre-washed 40 uL of packed M2 FLAG beads (Sigma) rotating at 4°C for 4 hrs. Afterwards, beads were washed with NP40 lysis buffer for a total of 4 times, 3 min each at 4°C. Before the last wash 20% of the IP was taken out and saved for western blot if needed. After the final wash with lysis buffer beads were transferred to a new tube and finally washed with 1x PBS (mixing by hand) and stored at – 80°C until mass spectrometry.

Mass spectrometry was performed by the proteomics resource facility at the Department of Pathology, University of Michigan. In brief, the beads were resuspended in 50 μl of 0.1M ammonium bicarbonate buffer (pH~8). Cysteines were reduced by adding 50 μl of 10 mM DTT and incubating at 45° C for 30 min. An overnight digestion with 1 ug sequencing grade, modified trypsin was carried out at 37° C with constant shaking in a Thermomixer. Samples were completely dried using vacufuge. Resulting peptides were dissolved in 8 μl of 0.1% formic acid/2% acetonitrile solution and 2 μls of the peptide solution were resolved on a nano-capillary reverse phase column (Acclaim PepMap C18, 2 micron, 50 cm, ThermoScientific) using a 0.1% formic acid/2% acetonitrile (Buffer A) and 0.1% formic acid/95% acetonitrile (Buffer B) gradient at 300 nl/min over a period of 180 min (2-22% buffer B in 110 min, 22-40% in 25 min, 40-90% in 5 min followed by holding at 90% buffer B for 5 min and requilibration with Buffer A for 25 min). Eluent was directly introduced into Orbitrap Fusion tribrid mass spectrometer (Thermo Scientific, San Jose CA) using an EasySpray source. Proteins were identified by searching the MS/MS data against *H Sapien* (UniProt; 20,145 reviewed entries; downloaded on 08-02-2017) using Proteome Discoverer (v2.1, Thermo Scientific). False discovery rate (FDR) was determined using Percolator and proteins/peptides with a FDR of ≤1% were retained for further analysis.

### 5. Drosophila lines and rough eye screening and survival assays

All fly lines used here and their sources are listed in **Supplementary Table 2**. To make the dSF2 OE fly line, *Drosophila* dSF2 sequence was PCR amplified (dSF2 F 5’-CACCATGGGATCACGCAACGAGTGCCG-3’ and dSF2 R 5’- ATAGTTAGAACGTGAGCGAGACCTGG-3’) was cloned into pEntry-TOPO vector (Thermo Fisher Scientific). The pENTR-dSF2 vector was recombined with Gateway plasmid pTWH (Drosophila Genomics Resource Center, IN). The final vector was used for site-specific transgenesis using PhiC31 integrase technique (BestGene, CA).

Flies were crossed and raised at 25°C on SY10 food supplemented with dry yeast unless otherwise noted. For rough eye screening, 5-6 virgin female flies expressing GMR-GAL4 driven UAS-*FMR1* (CGG)_90_◻EGFP reporters were crossed with male flies carrying either UAS◻driven siRNA against a candidate gene or a germline mutation. For GGGGCC repeat RNA toxicity modifier phenotyping a GMR-GAL4 driven UAS-(GGGGCC)_28_-EGFP reporter containing fly was used. Rough eye phenotypes in F_1_ progenies were scored at 1–2 days post-eclosion. A minimum of 30 flies (both male and females) from two independent crosses was scored. For rough eye scores were given based on following eye abnormalities: i. abnormal orientation of the bristles, ii. supernumerary bristles, iii. ommatidial fusion and disarray, iv. presence of necrosis and v. collapse/shrinkage of the eye. For each category three possible scores were given: 1 (for presence of the abnormality), 3 (if the abnormality affected >5% of the total eye) and 5 (if the abnormality affected >50% of the total eye). Eye images were captured using a Leica M125 stereomicroscope and a Leica DFC425 digital camera.

For survival assays, flies carrying desirable repeat RNA reporter and either a Tub5◻GAL4 GeneSwitch or ElaV-GAL4 GeneSwitch driver were crossed with a modifier fly. F_1_ progenies were collected 1 day post-eclosion and placed on SY10 food supplemented with 200 μM RU486 and flipped onto fresh RU486-containing food every 48 h. For survival, ~20 flies (equal male and females) from at least three independent crosses maintained at 29°C. Number of deaths recorded every 48 h until expiration and plotted using GraphPad Prism.

### 6. Primary rat neuron drug treatment, transfection and automated fluorescence microscopy imaging

Rat embryonic cortical dissections from E20 Long–Evans rat pups of both sexes were performed as previously described (Flores *et al*, 2019; Malik *et al*, 2018). Dissociated cortical neurons were plated at 0.6 × 10^5^ cells per well on poly-D-lysine coated 96-well plate in neuronal growth media (NGM, neurobasal A media, 2% B-27, 1% Glutamax-1 (v:v)), and maintained at 37°C for 4 days before transfection. On DIV 4, neurons were treated with SRPIN (10-50μM) or SPHINX (2-10μM) or DMSO 8 hours before transfection. Neurons were then cotransfected 0.1 μg of pGW1-mCherry and either 0.1 μg of pGW1-GFP or 0.1 μg of pGW +1(CGG)100 GFP DNA per well of a 96-well culture plate, using Lipofectamine 2000 (Thermo Fisher Scientific).

24 hours after transfection, neurons were imaged at 24 hour intervals for 10 days using an automated fluorescence microscopy platform previously described (Barmada *et al*, 2015; Arrasate *et al*, 2004). Images were processed using a custom code written in Python and ImageJ macro language, and analyzed by cox proportional hazard test using the survival package in R.

### 7. HCR and RNA foci analysis

HCR was performed as previously described (Rodriguez *et al*, 2020; Glineburg *et al*, 2021). Briefly, HEK293T cells were seeded at 1×10^5^ cells/mL in the chamber pre-coated with poly-D-lysine at the bottom. After transfection, cells were washed with 1xPBS twice then fixed in 4% PFA for 10 min at room temperature (RT). Fixative was aspirated and cells were washed with 1xPBS twice before treating with Turbo DNAse for 15 min at 37°C incubator. After washed with 1x PBS again, cells were dehydrated from 70% ethanol at 4°C overnight. Next, cells were quickly washed with 1x PBS then rehydrated in 1x PBS for 1 hr at RT. The ICC was done after fixing and before HCR. For ICC, cells were permeabilized in 0.1% TritonX-100 in 1x PBS for 6 min and block with 2% RNAse-free acetylated BSA in 1x PBS for 20 min at RT. Cells were stained with primary antibody SRSF1 (1:100, rabbit from Proteintech, 12929-2-AP) overnight at 4°C then followed by 3 times of 5 min 1x PBS washes before staining with secondary antibody (1:500, Alexa Fluor 488 goat anti-rabbit IgG from Invitrogen, A11008) for 1 hr at RT in dark. Then the cells was fixed again with 4% PFA for 10 min at RT followed by 3 times of 1 min 1x PBS washes before proceeding HCR. We follow HCRv2.0 protocol provided by Molecular Instrument. The initiator probe of (GGC)8 and fluorophore 647 labeled hairpin probes (B1H1 and B1H2) were synthesized by Molecular Instruments. Repeat RNA were detected by initiator probe (GGC)8 at 2 nM and amplified by fluorophore 647 labeled hairpin probes (B1H1 and B1H2) at 30 nM. Finally, cells were stained with DAPI, coverslipped in Prolong gold, and stored at 4°C in dark until imaging.

Fixed cells imaging was performed using a oil 60x objective in Olympus FV1000 inverted laser-scanning confocal microscope. For all experiments, acquisition parameters were identical between conditions within experiments. Cells were imaged in a series of Z-planes and images were analyzed in ImageJ. Average intensity composite images were derived from raw image files. For nucleus to cytoplasm ratio analysis, signals for each channel were normalized prior to quantification. The background signal was first normalized to non-transfection group or each corresponding control. Next, the ROI was applied to the DAPI channels to specify the region of nucleus along with the 647 channel which captured CGG RNA amplified by HCR. CGG RNA signal from the nucleus were calculated by the intensity in ROI from DAPI channel in pixels while the RNA signal from the cytoplasm were calculated by the total intensity from 647 channel subtracted by the RNA intensity in ROI from DAPI channel in pixels. Finally, nucleus to cytoplasm ratio was calculated by dividing the signal of CGG RNA from nucleus to cytoplasm. More than 588 of HEK293T cells were counted for each condition. P-values were calculated using two-tailed Student’s t-test with Bonferroni and Welch’s correction.

### 8. Statistical methods

Statistical analysis was performed using GraphPad Prism 7. For comparison of nLuc reporter luciferase activity, two-tailed Student’s t tests were performed with Welch’s correction. Two-tailed Student’s t-test with Bonferroni and Welch’s correction tests were used for quantifying HCR experiments, to determine if distribution of RNA and RNA foci in nucleus vs cytoplasm is statically significant between experimental groups.

### 9. Data availability

Raw mass spectrometry data is provided in Extended data Table.

## Acknowledgements

The authors thank Todd lab members for helpful input and instruction on Drosophila biology. We thank Amy Krans for assistance with Drosophila generation. We thank the Pletcher lab at the University of Michigan for providing all fly food. We acknowledge the proteomics resource facility at the Department of Pathology, University of Michigan.

## Reference

Anczuków O, Akerman M, Cléry A, Wu J, Shen C, Shirole NH, Raimer A, Sun S, Jensen MA, Hua Y, et al (2015) SRSF1-Regulated Alternative Splicing in Breast Cancer. Mol Cell 60: 105–117

Anczuków O, Rosenberg AZ, Akerman M, Das S, Zhan L, Karni R, Muthuswamy SK & Krainer AR (2012) The splicing factor SRSF1 regulates apoptosis and proliferation to promote mammary epithelial cell transformation. Nat Struct Mol Biol 19: 220–228

Ariza J, Rogers H, Monterrubio A, Reyes-Miranda A, Hagerman PJ & Martínez-Cerdeño V (2016) A Majority of FXTAS Cases Present with Intranuclear Inclusions Within Purkinje Cells. Cerebellum 15: 546–551

Arrasate M, Mitra S, Schweitzer ES, Segal MR & Finkbeiner S (2004) Inclusion body formation reduces levels of mutant huntingtin and the risk of neuronal death. Nature 431: 805–810

Ash PEA, Bieniek KF, Gendron TF, Caulfield T, Lin W-L, Dejesus-Hernandez M, van Blitterswijk MM, Jansen-West K, Paul JW, Rademakers R, et al (2013) Unconventional translation of C9ORF72 GGGGCC expansion generates insoluble polypeptides specific to c9FTD/ALS. Neuron 77: 639–646

Ayhan F, Perez BA, Shorrock HK, Zu T, Banez-Coronel M, Reid T, Furuya H, Clark HB, Troncoso JC, Ross CA, et al (2018) SCA8 RAN polySer protein preferentially accumulates in white matter regions and is regulated by eIF3F. EMBO J 37

Bañez-Coronel M, Ayhan F, Tarabochia AD, Zu T, Perez BA, Tusi SK, Pletnikova O, Borchelt DR, Ross CA, Margolis RL, et al (2015) RAN Translation in Huntington Disease. Neuron 88: 667–677

Barmada SJ, Ju S, Arjun A, Batarse A, Archbold HC, Peisach D, Li X, Zhang Y, Tank EMH, Qiu H, et al (2015) Amelioration of toxicity in neuronal models of amyotrophic lateral sclerosis by hUPF1. Proc Natl Acad Sci U S A 112: 7821–7826

Batson J, Toop HD, Redondo C, Babaei-Jadidi R, Chaikuad A, Wearmouth SF, Gibbons B, Allen C, Tallant C, Zhang J, et al (2017) Development of Potent, Selective SRPK1 Inhibitors as Potential Topical Therapeutics for Neovascular Eye Disease. ACS Chem Biol 12: 825–832

Bond U (2006) Stressed out! Effects of environmental stress on mRNA metabolism. FEMS Yeast Res 6: 160–170

Buijsen RAM, Sellier C, Severijnen L-AWFM, Oulad-Abdelghani M, Verhagen RFM, Berman RF, Charlet-Berguerand N, Willemsen R & Hukema RK (2014) FMRpolyG-positive inclusions in CNS and non-CNS organs of a fragile X premutation carrier with fragile X-associated tremor/ataxia syndrome. Acta Neuropathol Commun 2: 162

Chandra A, Ananda H, Singh N & Qamar I (2020) Identification of a novel and potent small molecule inhibitor of SRPK1: mechanism of dual inhibition of SRPK1 for the inhibition of cancer progression. Aging (Albany NY) 12

Chao JA, Patskovsky Y, Almo SC & Singer RH (2008) Structural basis for the coevolution of a viral RNA–protein complex. Nature Structural & Molecular Biology 15: 103–105

Cheng W, Wang S, Mestre AA, Fu C, Makarem A, Xian F, Hayes LR, Lopez-Gonzalez R, Drenner K, Jiang J, et al (2018) C9ORF72 GGGGCC repeat-associated non-AUG translation is upregulated by stress through eIF2α phosphorylation. Nature Communications 9: 51

Cheng W, Wang S, Zhang Z, Morgens DW, Hayes LR, Lee S, Portz B, Xie Y, Nguyen BV, Haney MS, et al (2019) CRISPR-Cas9 Screens Identify the RNA Helicase DDX3X as a Repressor of C9ORF72 (GGGGCC)n Repeat-Associated Non-AUG Translation. Neuron 104: 885–898.e8

Choi J-H, Kim S-H, Jeong Y-H, Kim SW, Min K-T & Kim K-T (2019) hnRNP Q Regulates Internal Ribosome Entry Site-Mediated fmr1 Translation in Neurons. Mol Cell Biol 39

Ciesiolka A, Jazurek M, Drazkowska K & Krzyzosiak WJ (2017) Structural Characteristics of Simple RNA Repeats Associated with Disease and their Deleterious Protein Interactions. Front Cell Neurosci 11: 97

Conlon EG, Lu L, Sharma A, Yamazaki T, Tang T, Shneider NA & Manley JL (2016) The C9ORF72 GGGGCC expansion forms RNA G-quadruplex inclusions and sequesters hnRNP H to disrupt splicing in ALS brains. Elife 5

Cooper-Knock J, Higginbottom A, Stopford MJ, Highley JR, Ince PG, Wharton SB, Pickering-Brown S, Kirby J, Hautbergue GM & Shaw PJ (2015) Antisense RNA foci in the motor neurons of C9ORF72-ALS patients are associated with TDP-43 proteinopathy. Acta Neuropathol 130: 63–75

Cooper-Knock J, Walsh MJ, Higginbottom A, Robin Highley J, Dickman MJ, Edbauer D, Ince PG, Wharton SB, Wilson SA, Kirby J, et al (2014) Sequestration of multiple RNA recognition motif-containing proteins by C9orf72 repeat expansions. Brain 137: 2040–2051

Coulon A, Ferguson ML, de Turris V, Palangat M, Chow CC & Larson DR (2014) Kinetic competition during the transcription cycle results in stochastic RNA processing. Elife 3

Dennis G, Sherman BT, Hosack DA, Yang J, Gao W, Lane HC & Lempicki RA (2003) DAVID: Database for Annotation, Visualization, and Integrated Discovery. Genome Biology 4: R60

Donnelly CJ, Zhang P-W, Pham JT, Haeusler AR, Heusler AR, Mistry NA, Vidensky S, Daley EL, Poth EM, Hoover B, et al (2013) RNA toxicity from the ALS/FTD C9ORF72 expansion is mitigated by antisense intervention. Neuron 80: 415–428

Fay MM, Anderson PJ & Ivanov P (2017) ALS/FTD-Associated C9ORF72 Repeat RNA Promotes Phase Transitions In Vitro and in Cells. Cell Rep 21: 3573–3584

Flores BN, Li X, Malik AM, Martinez J, Beg AA & Barmada SJ (2019) An Intramolecular Salt Bridge Linking TDP43 RNA Binding, Protein Stability, and TDP43-Dependent Neurodegeneration. Cell Reports 27: 1133–1150.e8

Fukuhara T, Hosoya T, Shimizu S, Sumi K, Oshiro T, Yoshinaka Y, Suzuki M, Yamamoto N, Herzenberg LA, Herzenberg LA, et al (2006) Utilization of host SR protein kinases and RNA-splicing machinery during viral replication. PNAS 103: 11329–11333

Gammons MV, Fedorov O, Ivison D, Du C, Clark T, Hopkins C, Hagiwara M, Dick AD, Cox R, Harper SJ, et al (2013) Topical antiangiogenic SRPK1 inhibitors reduce choroidal neovascularization in rodent models of exudative AMD. Invest Ophthalmol Vis Sci 54: 6052–6062

Gendron TF, Bieniek KF, Zhang Y-J, Jansen-West K, Ash PEA, Caulfield T, Daughrity L, Dunmore JH, Castanedes-Casey M, Chew J, et al (2013) Antisense transcripts of the expanded C9ORF72 hexanucleotide repeat form nuclear RNA foci and undergo repeat-associated non-ATG translation in c9FTD/ALS. Acta Neuropathol 126: 829–844

Glineburg MR, Todd PK, Charlet-Berguerand N & Sellier C (2018) Repeat-associated non-AUG (RAN) translation and other molecular mechanisms in Fragile X Tremor Ataxia Syndrome. Brain Res 1693: 43–54

Glineburg MR, Zhang Y, Tank EM, Barmada S & Todd P (2021) Enhanced detection of nucleotide repeat mRNA with hybridization chain reaction. bioRxiv: 2021.01.06.425640

Greco CM, Berman RF, Martin RM, Tassone F, Schwartz PH, Chang A, Trapp BD, Iwahashi C, Brunberg J, Grigsby J, et al (2006) Neuropathology of fragile X-associated tremor/ataxia syndrome (FXTAS). Brain 129: 243–255

Greco CM, Hagerman RJ, Tassone F, Chudley AE, Del Bigio MR, Jacquemont S, Leehey M & Hagerman PJ (2002) Neuronal intranuclear inclusions in a new cerebellar tremor/ataxia syndrome among fragile X carriers. Brain 125: 1760–1771

Green KM, Glineburg MR, Kearse MG, Flores BN, Linsalata AE, Fedak SJ, Goldstrohm AC, Barmada SJ & Todd PK (2017) RAN translation at C9orf72-associated repeat expansions is selectively enhanced by the integrated stress response. Nat Commun 8: 2005

Hagerman RJ, Leehey M, Heinrichs W, Tassone F, Wilson R, Hills J, Grigsby J, Gage B & Hagerman PJ (2001) Intention tremor, parkinsonism, and generalized brain atrophy in male carriers of fragile X. Neurology 57: 127–130

Harlen KM & Churchman LS (2017) Subgenic Pol II interactomes identify region-specific transcription elongation regulators. Mol Syst Biol 13: 900

Hautbergue GM, Castelli LM, Ferraiuolo L, Sanchez-Martinez A, Cooper-Knock J, Higginbottom A, Lin Y-H, Bauer CS, Dodd JE, Myszczynska MA, et al (2017) SRSF1-dependent nuclear export inhibition of C9ORF72 repeat transcripts prevents neurodegeneration and associated motor deficits. Nat Commun 8

He F, Flores BN, Krans A, Frazer M, Natla S, Niraula S, Adefioye O, Barmada SJ & Todd PK (2020) The carboxyl termini of RAN translated GGGGCC nucleotide repeat expansions modulate toxicity in models of ALS/FTD. Acta Neuropathol Commun 8: 122

Hinnebusch AG (2006) eIF3: a versatile scaffold for translation initiation complexes. Trends Biochem Sci 31: 553–562

Hong Y, Chan CB, Kwon I-S, Li X, Song M, Lee H-P, Liu X, Sompol P, Jin P, Lee H, et al (2012) SRPK2 Phosphorylates Tau and Mediates the Cognitive Defects in Alzheimer’s Disease. J Neurosci 32: 17262–17272

Ishiguro T, Sato N, Ueyama M, Fujikake N, Sellier C, Kanegami A, Tokuda E, Zamiri B, Gall-Duncan T, Mirceta M, et al (2017) Regulatory Role of RNA Chaperone TDP-43 for RNA Misfolding and Repeat-Associated Translation in SCA31. Neuron 94: 108–124.e7

Jacquemont S, Hagerman RJ, Leehey M, Grigsby J, Zhang L, Brunberg JA, Greco C, Des Portes V, Jardini T, Levine R, et al (2003) Fragile X Premutation Tremor/Ataxia Syndrome: Molecular, Clinical, and Neuroimaging Correlates. The American Journal of Human Genetics 72: 869–878

Jacquemont S, Hagerman RJ, Leehey MA, Hall DA, Levine RA, Brunberg JA, Zhang L, Jardini T, Gane LW, Harris SW, et al (2004) Penetrance of the fragile X-associated tremor/ataxia syndrome in a premutation carrier population. JAMA 291: 460–469

Jain A & Vale RD (2017) RNA phase transitions in repeat expansion disorders. Nature 546: 243–247

Jazurek M, Ciesiolka A, Starega-Roslan J, Bilinska K & Krzyzosiak WJ (2016) Identifying proteins that bind to specific RNAs - focus on simple repeat expansion diseases. Nucleic Acids Res 44: 9050–9070

Jeong S (2017) SR Proteins: Binders, Regulators, and Connectors of RNA. Mol Cells 40: 1–9

Jin P, Duan R, Qurashi A, Qin Y, Tian D, Rosser TC, Liu H, Feng Y & Warren ST (2007) Pur α binds to rCGG repeats and modulates repeat-mediated neurodegeneration in a Drosophila model of Fragile X Tremor/Ataxia Syndrome. Neuron 55: 556–564

Jovičić A, Mertens J, Boeynaems S, Bogaert E, Chai N, Yamada SB, Paul JW, Sun S, Herdy JR, Bieri G, et al (2015) Modifiers of C9orf72 dipeptide repeat toxicity connect nucleocytoplasmic transport defects to FTD/ALS. Nat Neurosci 18: 1226–1229

Kanadia RN, Johnstone KA, Mankodi A, Lungu C, Thornton CA, Esson D, Timmers AM, Hauswirth WW & Swanson MS (2003) A muscleblind knockout model for myotonic dystrophy. Science 302: 1978–1980

Kanadia RN, Shin J, Yuan Y, Beattie SG, Wheeler TM, Thornton CA & Swanson MS (2006) Reversal of RNA missplicing and myotonia after muscleblind overexpression in a mouse poly(CUG) model for myotonic dystrophy. PNAS 103: 11748–11753

Karni R, de Stanchina E, Lowe SW, Sinha R, Mu D & Krainer AR (2007) The gene encoding the splicing factor SF2/ASF is a proto-oncogene. Nat Struct Mol Biol 14: 185–193

Kearse MG, Green KM, Krans A, Rodriguez CM, Linsalata AE, Goldstrohm AC & Todd PK (2016) CGG Repeat-Associated Non-AUG Translation Utilizes a Cap-Dependent Scanning Mechanism of Initiation to Produce Toxic Proteins. Mol Cell 62: 314–322

Krans A, Kearse MG & Todd PK (2016) Repeat-associated non-AUG translation from antisense CCG repeats in fragile X tremor/ataxia syndrome. Ann Neurol 80: 871–881

Krzyzosiak WJ, Sobczak K, Wojciechowska M, Fiszer A, Mykowska A & Kozlowski P (2012) Triplet repeat RNA structure and its role as pathogenic agent and therapeutic target. Nucleic Acids Res 40: 11–26

Lee K-H, Zhang P, Kim HJ, Mitrea DM, Sarkar M, Freibaum BD, Cika J, Coughlin M, Messing J, Molliex A, et al (2016) C9orf72 Dipeptide Repeats Impair the Assembly, Dynamics, and Function of Membrane-Less Organelles. Cell 167: 774–788.e17

Lee Y-B, Chen H-J, Peres JN, Gomez-Deza J, Attig J, Stalekar M, Troakes C, Nishimura AL, Scotter EL, Vance C, et al (2013) Hexanucleotide repeats in ALS/FTD form length-dependent RNA foci, sequester RNA binding proteins, and are neurotoxic. Cell Rep 5: 1178–1186

Linsalata AE, He F, Malik AM, Glineburg MR, Green KM, Natla S, Flores BN, Krans A, Archbold HC, Fedak SJ, et al (2019) DDX3X and specific initiation factors modulate FMR1 repeat-associated non-AUG-initiated translation. EMBO Rep 20: e47498

Maimon A, Mogilevsky M, Shilo A, Golan-Gerstl R, Obiedat A, Ben-Hur V, Lebenthal-Loinger I, Stein I, Reich R, Beenstock J, et al (2014) Mnk2 alternative splicing modulates the p38-MAPK pathway and impacts Ras-induced transformation. Cell Rep 7: 501–513

Malik AM, Miguez RA, Li X, Ho Y-S, Feldman EL & Barmada SJ (2018) Matrin 3-dependent neurotoxicity is modified by nucleic acid binding and nucleocytoplasmic localization. Elife 7

Mankodi A, Logigian E, Callahan L, McClain C, White R, Henderson D, Krym M & Thornton CA (2000) Myotonic dystrophy in transgenic mice expressing an expanded CUG repeat. Science 289: 1769–1773

Matheny T, Treeck BV, Huynh TN & Parker R (2020) RNA partitioning into stress granules is based on the summation of multiple interactions. RNA: rna.078204.120

May S, Hornburg D, Schludi MH, Arzberger T, Rentzsch K, Schwenk BM, Grässer FA, Mori K, Kremmer E, Banzhaf-Strathmann J, et al (2014) C9orf72 FTLD/ALS-associated Gly-Ala dipeptide repeat proteins cause neuronal toxicity and Unc119 sequestration. Acta Neuropathol 128: 485–503

Miller JW, Urbinati CR, Teng-Umnuay P, Stenberg MG, Byrne BJ, Thornton CA & Swanson MS (2000) Recruitment of human muscleblind proteins to (CUG)(n) expansions associated with myotonic dystrophy. EMBO J 19: 4439–4448

Mizielinska S, Grönke S, Niccoli T, Ridler CE, Clayton EL, Devoy A, Moens T, Norona FE, Woollacott IOC, Pietrzyk J, et al (2014) C9orf72 repeat expansions cause neurodegeneration in Drosophila through arginine-rich proteins. Science 345: 1192–1194

Mori K, Weng S-M, Arzberger T, May S, Rentzsch K, Kremmer E, Schmid B, Kretzschmar HA, Cruts M, Van Broeckhoven C, et al (2013) The C9orf72 GGGGCC repeat is translated into aggregating dipeptide-repeat proteins in FTLD/ALS. Science 339: 1335–1338

Nakagawa O, Arnold M, Nakagawa M, Hamada H, Shelton JM, Kusano H, Harris TM, Childs G, Campbell KP, Richardson JA, et al (2005) Centronuclear myopathy in mice lacking a novel muscle-specific protein kinase transcriptionally regulated by MEF2. Genes Dev 19: 2066–2077

Ong S-E, Blagoev B, Kratchmarova I, Kristensen DB, Steen H, Pandey A & Mann M (2002) Stable isotope labeling by amino acids in cell culture, SILAC, as a simple and accurate approach to expression proteomics. Mol Cell Proteomics 1: 376–386

Prudencio M, Belzil VV, Batra R, Ross CA, Gendron TF, Pregent LJ, Murray ME, Overstreet KK, Piazza-Johnston AE, Desaro P, et al (2015) Distinct brain transcriptome profiles in C9orf72-associated and sporadic ALS. Nature Neuroscience 18: 1175–1182

Rizzotto D, Zaccara S, Rossi A, Galbraith MD, Andrysik Z, Pandey A, Sullivan KD, Quattrone A, Espinosa JM, Dassi E, et al (2020) Nutlin-Induced Apoptosis Is Specified by a Translation Program Regulated by PCBP2 and DHX30. Cell Rep 30: 4355–4369.e6

Rodriguez CM, Wright SE, Kearse MG, Haenfler JM, Flores BN, Liu Y, Ifrim MF, Glineburg MR, Krans A, Jafar-Nejad P, et al (2020) A native function for RAN translation and CGG repeats in regulating fragile X protein synthesis. Nat Neurosci 23: 386–397

van Roosmalen W, Le Dévédec SE, Golani O, Smid M, Pulyakhina I, Timmermans AM, Look MP, Zi D, Pont C, de Graauw M, et al (2015) Tumor cell migration screen identifies SRPK1 as breast cancer metastasis determinant. J Clin Invest 125: 1648–1664

Sato N, Amino T, Kobayashi K, Asakawa S, Ishiguro T, Tsunemi T, Takahashi M, Matsuura T, Flanigan KM, Iwasaki S, et al (2009) Spinocerebellar ataxia type 31 is associated with ‘inserted’ penta-nucleotide repeats containing (TGGAA)n. Am J Hum Genet 85: 544–557

Sellier C, Buijsen RAM, He F, Natla S, Jung L, Tropel P, Gaucherot A, Jacobs H, Meziane H, Vincent A, et al (2017) Translation of Expanded CGG Repeats into FMRpolyG Is Pathogenic and May Contribute to Fragile X Tremor Ataxia Syndrome. Neuron 93: 331–347

Sellier C, Freyermuth F, Tabet R, Tran T, He F, Ruffenach F, Alunni V, Moine H, Thibault C, Page A, et al (2013) Sequestration of DROSHA and DGCR8 by expanded CGG RNA repeats alters microRNA processing in fragile X-associated tremor/ataxia syndrome. Cell Rep 3: 869–880

Sellier C, Rau F, Liu Y, Tassone F, Hukema RK, Gattoni R, Schneider A, Richard S, Willemsen R, Elliott DJ, et al (2010) Sam68 sequestration and partial loss of function are associated with splicing alterations in FXTAS patients. EMBO J 29: 1248–1261

Sheng J, Zhao Q, Zhao J, Zhang W, Sun Y, Qin P, Lv Y, Bai L, Yang Q, Chen L, et al (2018) SRSF1 modulates PTPMT1 alternative splicing to regulate lung cancer cell radioresistance. EBioMedicine 38: 113–126

Sofola OA, Jin P, Qin Y, Duan R, Liu H, de Haro M, Nelson DL & Botas J (2007) RNA-binding proteins hnRNP A2/B1 and CUGBP1 suppress fragile X CGG premutation repeat-induced neurodegeneration in a Drosophila model of FXTAS. Neuron 55: 565–571

Sonobe Y, Ghadge G, Masaki K, Sendoel A, Fuchs E & Roos RP (2018) Translation of dipeptide repeat proteins from the C9ORF72 expanded repeat is associated with cellular stress. Neurobiol Dis 116: 155–165

Soragni E, Petrosyan L, Rinkoski TA, Wieben ED, Baratz KH, Fautsch MP & Gottesfeld JM (2018) Repeat-Associated Non-ATG (RAN) Translation in Fuchs’ Endothelial Corneal Dystrophy. Invest Ophthalmol Vis Sci 59: 1888–1896

Taneja KL, McCurrach M, Schalling M, Housman D & Singer RH (1995) Foci of trinucleotide repeat transcripts in nuclei of myotonic dystrophy cells and tissues. J Cell Biol 128: 995–1002

Tassone F, Iong KP, Tong T-H, Lo J, Gane LW, Berry-Kravis E, Nguyen D, Mu LY, Laffin J, Bailey DB, et al (2012) FMR1 CGG allele size and prevalence ascertained through newborn screening in the United States. Genome Med 4: 100

Todd PK, Oh SY, Krans A, He F, Sellier C, Frazer M, Renoux AJ, Chen K, Scaglione KM, Basrur V, et al (2013) CGG repeat-associated translation mediates neurodegeneration in fragile X tremor ataxia syndrome. Neuron 78: 440–455

Todd PK, Oh SY, Krans A, Pandey UB, Di Prospero NA, Min K-T, Taylor JP & Paulson HL (2010) Histone deacetylases suppress CGG repeat-induced neurodegeneration via transcriptional silencing in models of fragile X tremor ataxia syndrome. PLoS Genet 6: e1001240

Van Treeck B & Parker R (2018) Emerging Roles for Intermolecular RNA-RNA Interactions in RNP Assemblies. Cell 174: 791–802

Wang ET, Treacy D, Eichinger K, Struck A, Estabrook J, Olafson H, Wang TT, Bhatt K, Westbrook T, Sedehizadeh S, et al (2019) Transcriptome alterations in myotonic dystrophy skeletal muscle and heart. Hum Mol Genet 28: 1312–1321

Wang HY, Lin W, Dyck JA, Yeakley JM, Songyang Z, Cantley LC & Fu XD (1998) SRPK2: a differentially expressed SR protein-specific kinase involved in mediating the interaction and localization of pre-mRNA splicing factors in mammalian cells. J Cell Biol 140: 737–750

Wen X, Tan W, Westergard T, Krishnamurthy K, Markandaiah SS, Shi Y, Lin S, Shneider NA, Monaghan J, Pandey UB, et al (2014) Antisense proline-arginine RAN dipeptides linked to C9ORF72-ALS/FTD form toxic nuclear aggregates that initiate in vitro and in vivo neuronal death. Neuron 84: 1213–1225

Westergard T, McAvoy K, Russell K, Wen X, Pang Y, Morris B, Pasinelli P, Trotti D & Haeusler A (2019) Repeat-associated non-AUG translation in C9orf72-ALS/FTD is driven by neuronal excitation and stress. EMBO Mol Med 11

Yamada SB, Gendron TF, Niccoli T, Genuth NR, Grosely R, Shi Y, Glaria I, Kramer NJ, Nakayama L, Fang S, et al (2019) RPS25 is required for efficient RAN translation of C9orf72 and other neurodegenerative disease-associated nucleotide repeats. Nature Neuroscience 22: 1383–1388

Yan X, Hoek TA, Vale RD & Tanenbaum ME (2016) Dynamics of Translation of Single mRNA Molecules In Vivo. Cell 165: 976–989

Zhang Y-J, Gendron TF, Ebbert MTW, O’Raw AD, Yue M, Jansen-West K, Zhang X, Prudencio M, Chew J, Cook CN, et al (2018) Poly(GR) impairs protein translation and stress granule dynamics in C9orf72-associated frontotemporal dementia and amyotrophic lateral sclerosis. Nat Med 24: 1136–1142

Zhang Y-J, Guo L, Gonzales PK, Gendron TF, Wu Y, Jansen-West K, O’Raw AD, Pickles SR, Prudencio M, Carlomagno Y, et al (2019) Heterochromatin anomalies and double-stranded RNA accumulation underlie C9orf72 poly(PR) toxicity. Science 363

Zhou Z & Fu X-D (2013) Regulation of splicing by SR proteins and SR protein-specific kinases. Chromosoma 122: 191–207

Zu T, Cleary JD, Liu Y, Bañez-Coronel M, Bubenik JL, Ayhan F, Ashizawa T, Xia G, Clark HB, Yachnis AT, et al (2017) RAN Translation Regulated by Muscleblind Proteins in Myotonic Dystrophy Type 2. Neuron 95: 1292–1305.e5

Zu T, Gibbens B, Doty NS, Gomes-Pereira M, Huguet A, Stone MD, Margolis J, Peterson M, Markowski TW, Ingram MAC, et al (2011) Non-ATG-initiated translation directed by microsatellite expansions. Proc Natl Acad Sci U S A 108: 260–265

Zu T, Liu Y, Bañez-Coronel M, Reid T, Pletnikova O, Lewis J, Miller TM, Harms MB, Falchook AE, Subramony SH, et al (2013) RAN proteins and RNA foci from antisense transcripts in C9ORF72 ALS and frontotemporal dementia. Proc Natl Acad Sci U S A 110: E4968–4977

