## Supplemental files for "*In vivo* CGG repeat RNA binding protein capture identifies RAN translation modifiers and suppressors of repeat toxicity"

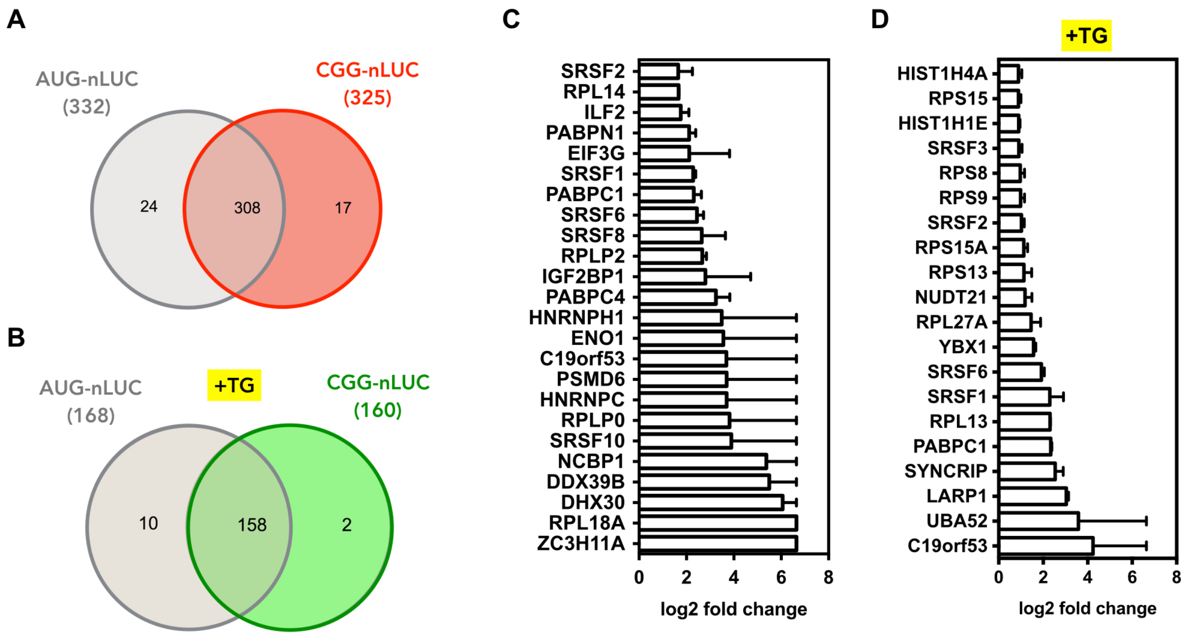


**Supplementary Figure 1.**

Venn diagrams indicate total number of proteins identified for AUG-nLUC and CGG-nLUC RNA-tagging reporters without (A) or with (B) TG treatment. List of top 20 proteins enriched in CGG-PP7 RNA interaction compared to AUG in mass spectrometry experiments without (C) or with (D) TG treatment. Bar graphs represent average of two biological replicates +/- range

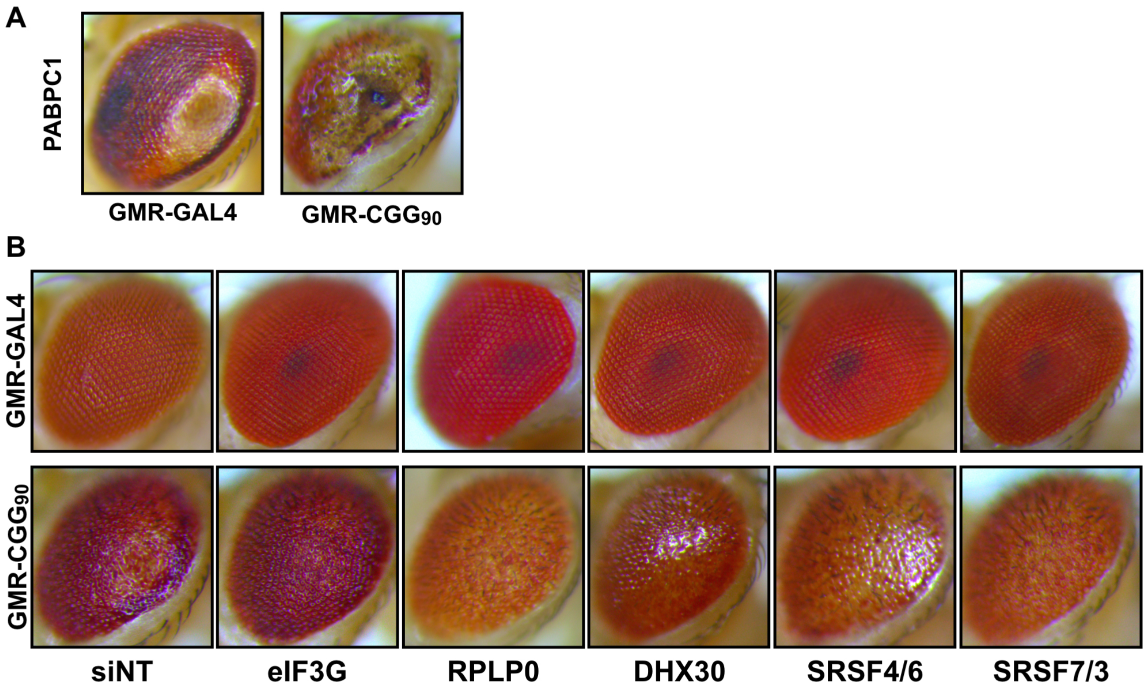


**Supplementary Figure 2.**

Representative photographs of fly eyes expressing either GMR-GAL4 driver alone or the (CGG)90-EGFP construct under a GMR-GAL4 driver, with siRNA against fly homologs of PABPC1 (A) and eIF3G, RPLP0, DHX30, SRSF4/6 and SRSF7/3 (B).

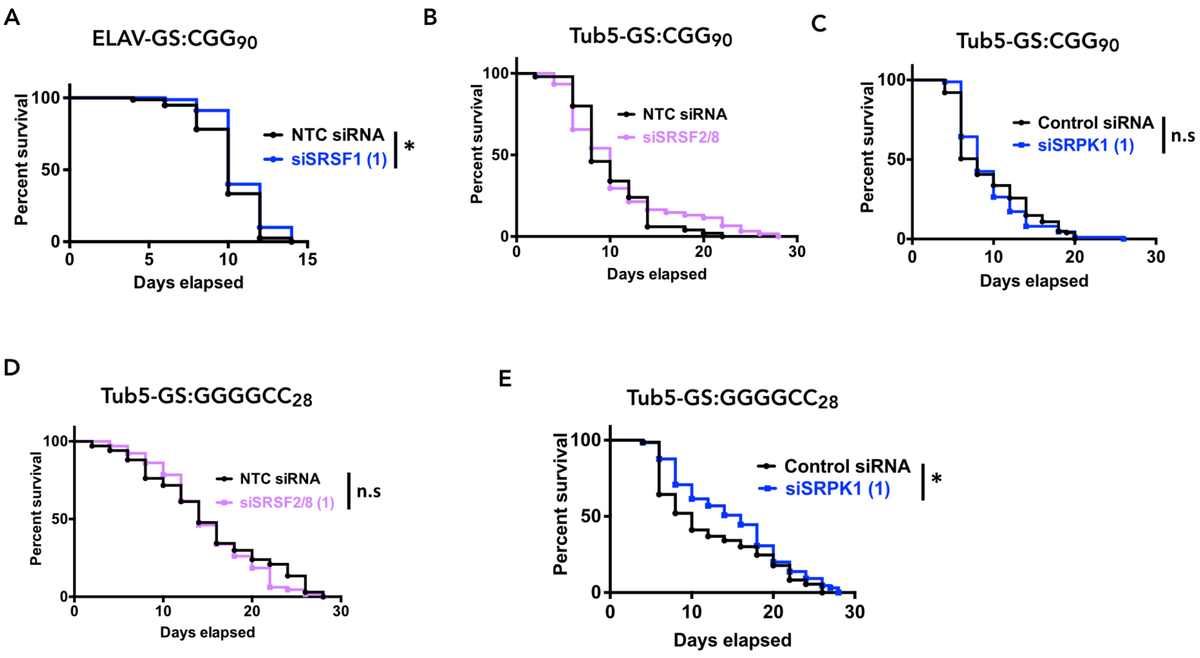


**Supplementary Figure 3.**

Survival assays of (CGG)90-EGFP and (G4C2)28-EGFP expressing fly under Tub5-GS or ELAV-GS drivers with respective siRNAs as mentioned (Log-rank Mantel–Cox test; n = 50-100/genotype). *$p < 0.05$

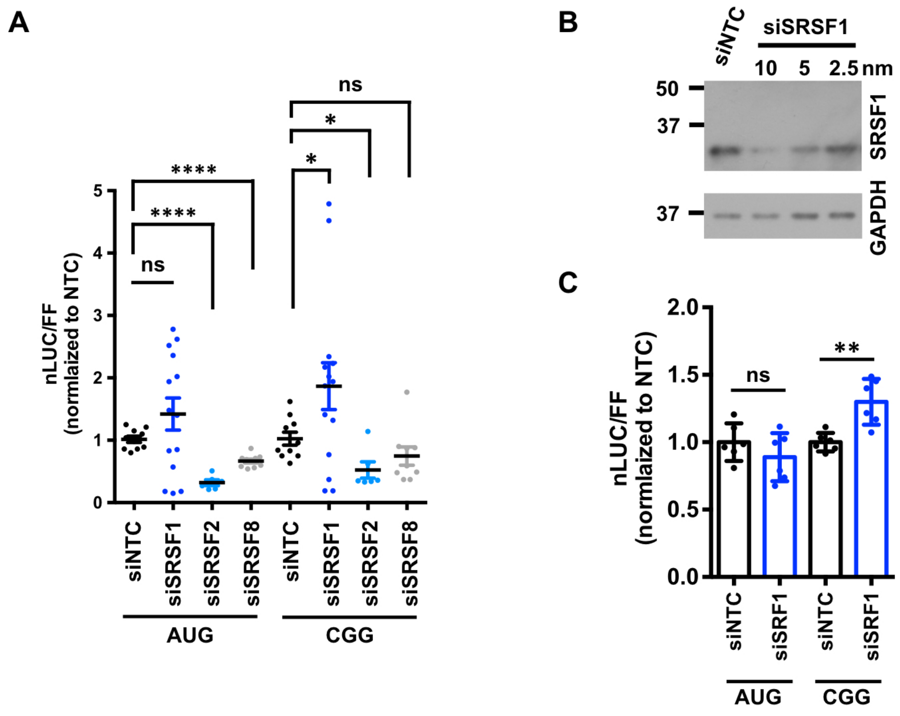


**Supplementary Figure 4**

(A) Relative expression of AUG-nLuc and CGG-nLuc reporters in HEK293T cells (n=6-14) following knockdown of SRSF1, SRSF2 and SRSF8. SRSF1 data points are same as presented in Figure 4B.

(B) Comparisons between siEGFP and siSRSF1 treated cells for confirmation of SRSF1 knockdown with a separate set of siRNA

(C) Relative expression of AUG-nLuc and CGG-nLuc reporters in HEK293T cells (n=6) following knockdown of SRSF1 with siRNA described in (B). t-test with Welch correction; *$p < 0.05$; **$p < 0.01$ and ****$p < 0.0001$

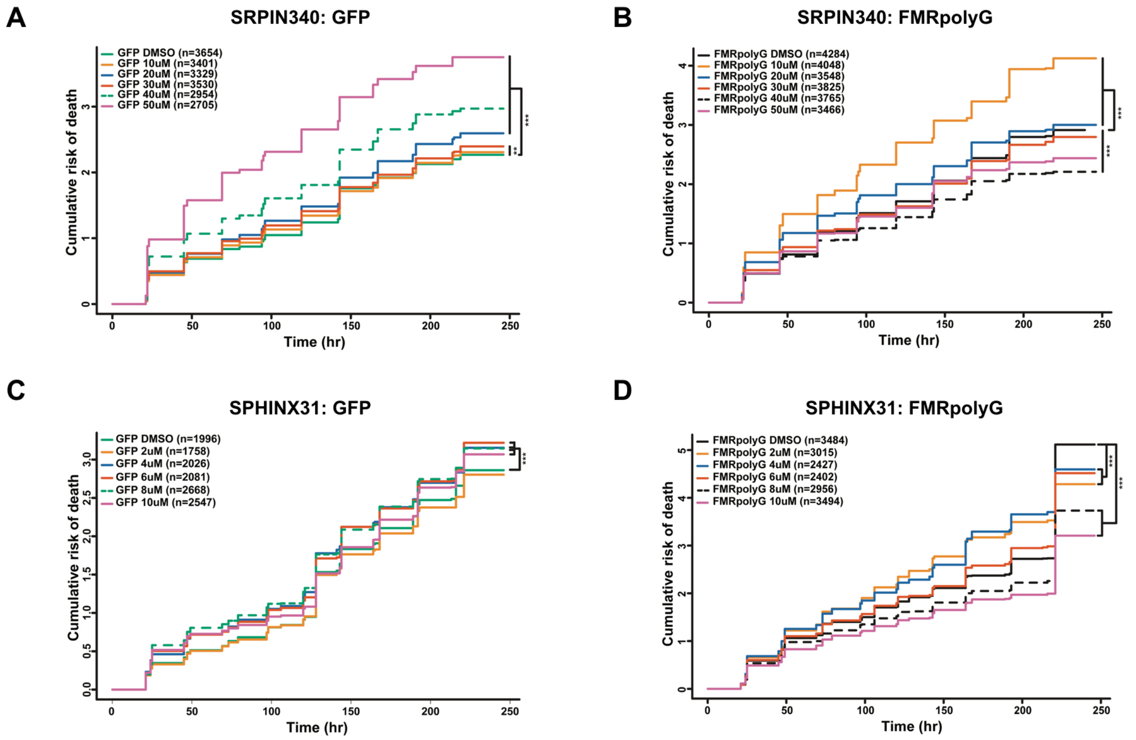


**Supplementary Figure 5.**

Pharmacological targeting of SPRK1 with range of concentrations of SRPIN340 (A-B) and SPHINX31 (C-D) showing the effects on GFP control (A and C) or +1(CGG)100-EGFP (B and D) expressing neurons.

**Supplementary Table 1**

Sequence maps of reporters used in this study

**AUG-nLUC-PP7:** T7-**ATG**nLUC-2xPP7

GAAATtaatacgactcactatagggAGACCCAAGCTGGCTAGCGTTTAAACTTAAGCTTGGCAATCCG
GTACTGTTGGTAAAGCCACC**ATG**gtcttcacactcgaagatttcgttggggactggcgacagacagccggctacaacct
ggaccaagtccttgaacagggaggtgtgtccagtttgtttcagaatctcggggtgtccgtaactccgatccaaaggattgtcctgagcggt
gaaaatgggctgaagatcgacatccatgtcatcatcccgtatgaaggtctgagcggcgaccaaatgggccagatcgaaaaaatttttaa
ggtggtgtaccctgtggatgatcatcactttaaggtgatcctgcactatggcacactggtaatcgacggggttacgccgaacatgatcgac
tatttcggacggccgtatgaaggcatcgccgtgttcgacggcaaaaagatcactgtaacagggaccctgtggaacggcaacaaaatta
tcgacgagcgcctgatcaaccccgacggctccctgctgttccgagtaaccatcaacggagtgaccggctggcggctgtgcgaacgcat
tctggcgTAAGGCCGCGACTCTAGAGggcccAGATTCACCGGTTAAGGTACCTAATTGCCTAGAA
AGGAGCAGACGATATGGCGTCGCTCCCTGCAGGTCGACTCTAGAAACCAGCAGAGCATATG
GGCTCGCTGGCTGCAGTATTCCCGGGTTCATTTAAGGTACCTAATTGCCTAGAAAGGAGCA
GACGATATGGCGTCGCTCCCTGCAGGTCGACTCTAGAAACCAGCAGAGCATATGGGCTCGC
TGGCTGCAGTATTCCCGGGTTCATTaccggt

**CGG-nLUC-PP7:** T7-FMR1 5'**-(CGG)90-GGG**nLUC-2xPP7

GAAATtaatacgactcactatagggAGACCCAAGCTGGCTAGCGTTTAAACTTAAGCTTGGTACCGAGC
TCGGATCCACTAGTCCAGTGTGGTGGAATTCGTTAACAGATCTGCTCAGCTCCGTTTCGGTT
TCacttccggtggagggccgcctctgagcgggcggcgggccgacggcgagcgcgggcggcggcggtgacggaggcgccgctgc
caggggcgtgcggcagcg**(CGG)90**CGCTGGGCCTCGAGGATATCAAGATCTGGCCTCGGCGGCC
AAGCTTGGCAATCCGGTACTGTTGGTAAAGCCACCC**GGG**gtcttcacactcgaagatttcgttggggactggc
gacagacagccggctacaacctggaccaagtccttgaacagggaggtgtgtccagtttgtttcagaatctcggggtgtccgtaactccga
tccaaaggattgtcctgagcggtgaaaatgggctgaagatcgacatccatgtcatcatcccgtatgaaggtctgagcggcgaccaaatg
ggccagatcgaaaaaatttttaaggtggtgtaccctgtggatgatcatcactttaaggtgatcctgcactatggcacactggtaatcgacgg
ggttacgccgaacatgatcgactatttcggacggccgtatgaaggcatcgccgtgttcgacggcaaaaagatcactgtaacagggacc
ctgtggaacggcaacaaaattatcgacgagcgcctgatcaaccccgacggctccctgctgttccgagtaaccatcaacggagtgaccg
gctggcggctgtgcgaacgcattctggcgTAAGGCCGCGACTCTAGAGggcccAGATTCACCGGTTAAGGTA
CCTAATTGCCTAGAAAGGAGCAGACGATATGGCGTCGCTCCCTGCAGGTCGACTCTAGAAA
CCAGCAGAGCATATGGGCTCGCTGGCTGCAGTATTCCCGGGTTCATTTAAGGTACCTAATTG
CCTAGAAAGGAGCAGACGATATGGCGTCGCTCCCTGCAGGTCGACTCTAGAAACCAGCAGA
GCATATGGGCTCGCTGGCTGCAGTATTCCCGGGTTCATTaccggt

**PCP-NLS-3xFLAG:** T7-**ATG**PCP-NLS-**3xFLAG**

GAAATaatacgactcactatagggAGACCCAAGCTGGCTAGCGTTTAAACTTAAGCTTGgtacCTCTCG
AGAATTCTCACGCGCCGGATCCGCCACC**ATG**TCCAAAACCATCGTTCTTTCGGTCGGCGAG
GCTACTCGCACTCTGACTGAGATCCAGTCCACCGCAGACCGTCAGATCTTCGAAGAGAAGG
TCGGGCCTCTGGTGGGTCGGCTGCGCCTCACGGCTTCGCTCCGTCAAAACGGAGCCAAGA
CCGCGTATCGCGTCAACCTAAAACTGGATCAGGCGGACGTCGTTGATTGCTCCACCAGCGT
CTGCGGCGAGCTTCCGAAAGTGCGCTACACTCAGGTATGGTCGCACGACGTGACAATCGTT
GCGAATAGCACCGAGGCCTCGCGCAAATCGTTGTACGATTTGACCAAGTCCCTCGTCGCGA
CCTCGCAGGTCGAAGATCTTGTCGTCAACCTTGTGCCGCTGGGCCGTCGTGCGGACCCGCT
AGCCTCCTGCGGCCGCCCAAAAAAGAagagaaaggtagaagaccccGACTACAAAGACCATGACGG
TGATTATAAAGATCATGACATCGATTACAAGGATGACGATGACAAGtaa

**Supplementary Table 2**
List of flies used in this study

| Modifier/target gene | Drosophila homolog | Source/stock number |
|---|---|---|
| HNRNPH | Glo | BDSC 33668 |
| PABPN1 | Pabp2 | BDSC 34602 |
| NCBP1 | Cbp80 | VDRC 110673 |
| EIF3G | eIF3G1 | BDSC 43243 |
| RPLP0 | RpLP0 | BDSC 31370 |
| LARP1 (1) | larp | BDSC 11687 |
| LARP1 (2) | larp | BDSC 42578 |
| DHX30 | bgcn | VDRC 108334 |
| DDX39B (1) | Hel25E | BDSC 11043 |
| DDX39B (2) | Hel25E | BDSC 33666 |
| SYNCRIP (1) | Syp | BDSC 55577 |
| SYNCRIP (2) | Syp | BDSC 56972 |
| SRSF1 (1) | SF2 | BDSC 32367 |
| SRSF1 (2) | SF2 | BDSC 29522 |
| SRSF2/8 (1) | SC35 | BDSC 65888 |
| SRSF2/8 (2) | SC35 | BDSC 20169 |
| SRSF4/6 (1) | B52 | BDSC 10265 |
| SRSF4/6 (2) | B52 | BDSC 37519 |
| SRSF7/3 (1) | x16 | BDSC 51468 |
| SRSF7/3 (2) | x16 | BDSC 55642 |
| SRPK1 (1) | SRPK | BDSC 57295 |
| SRPK1 (2) | SRPK | BDSC 57587 |
| Non-targeting Control | mCherry RNAi | BDSC 35785 |
| Non-targeting Control | LUC RNAi | BDSC 31603/Todd lab (G4C2/LUC) |
| Non-targeting Control | lexA RNAI | BDSC 67947/Todd lab (G4C2/lexA) |
| SRSF1 overexpression | SF2 | Todd lab |
